# Effect of molecular weight of tyramine-modified hyaluronan on polarization state of THP-1 and peripheral blood mononuclear cells-derived macrophages

**DOI:** 10.1101/2024.01.11.575241

**Authors:** Jacek K. Wychowaniec, Ezgi Irem Bektas, Andrea J. Vernengo, Marcia Mürner, Marielle Airoldi, Paul Sean Tipay, Jiranuwat Sapudom, Jeremy Teo, David Eglin, Matteo D’Este

## Abstract

The immunomodulatory properties of hyaluronan and its derivatives are key to their use in medicine and tissue engineering. In this work we evaluated the capability of soluble tyramine-modified hyaluronan (THA) of two molecular weights (low Mw=280 kDa and high Mw=1640 kDa) for polarization of THP-1 and peripheral blood mononuclear cells (PBMCs)-derived macrophages (MΦs). We demonstrate the polarization effects of the supplemented THA by flow cytometry and multiplex ELISA for the THP-1 derived MΦs and by semi-automated image analysis from confocal microscopy, immunofluorescent staining utilising CD68 and CD206 surface markers, RT-qPCR gene expression analysis, as well as using the enzyme-linked immunosorbent assay (ELISA) for PBMCs-derived MΦs. Our data indicate that supplementation with LMW THA drives changes in THP-1 derived MΦs towards a pro-inflammatory M1-like phenotype, whereas supplementation with the HMW THA leads to a more mixed profile with some features of both M1 and M2 phenotypes, suggesting either a heterogeneous population or a transitional state. For cells directly sourced from human patients, PMBCs-derived MΦs, results exhibit a higher degree of variability, pointing out a differential regulation of factors including IL-10 and CD206 between the two cell sources. While human primary cells add to the clinical relevance, donor diversity introduces wider variability in the dataset, preventing drawing strong conclusions. Nevertheless, the MΦs profiles observed in THP-1 derived cells for treatments with LMW and HMW THA are generally consistent with what might be expected for the treatment with non-modified hyaluronans of respective molecular weights, confirming the known association holds true for the chemically tyramine-modified hyaluronan. We stipulate that these responses will provide basis for more accurate in vivo representation and translational immunomodulatory guidance for the use of THA-based biomaterials to a wider biomaterials and tissue engineering communities.

**Graphical Abstract:** 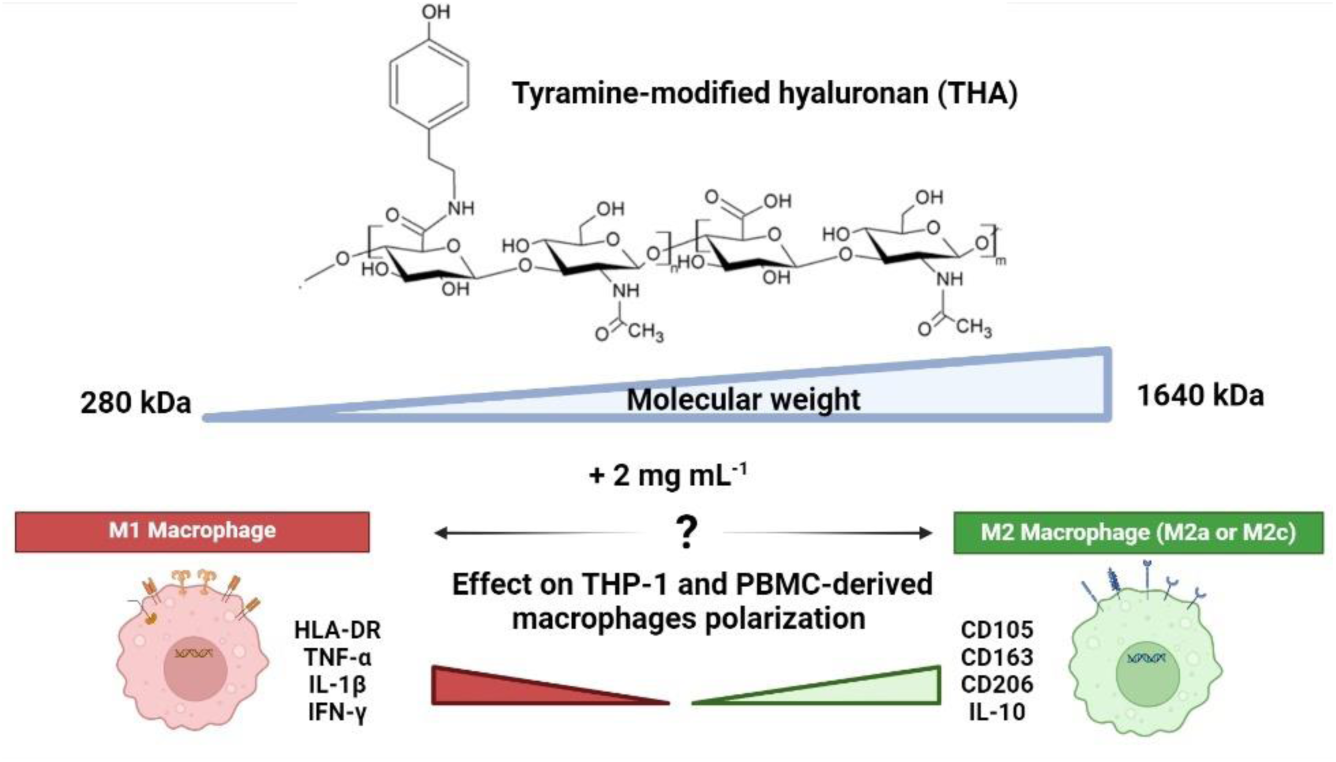

## 1. Introduction

The ability to generate and control a variety of individualized cellular signals plays a vital role in the designing biopolymers for biomedical applications. The immune system is essential for normal tissue healing, mediated by both innate and adaptive immunity, primarily through inflammation [1, 2]. During injury, the immune system can improve tissue repair or escalate degeneration, depending on the context [3, 4]. These effects are tied to the phenotypic state of the immune cells present at the injury site, particularly macrophages (MΦs). MΦs, among the most critical immune cells produce cytokines and chemokines that drive tissue repair or degeneration [5][6]. In the past decade, new MΦ subtypes have been identified, playing a key regulatory role in tissue regeneration [7]. MΦs thus exist on a spectrum, ranging from a pro-inflammatory M1 to anti-inflammatory M2 states [4, 8]. M1 MΦs respond to pathogens, produce pro-inflammatory cytokines (IL-1β, IL-6, TNF-α) and promote inflammation and host defence, whileM2 MΦs aid in tissue remodelling and healing in response to anti-inflammatory signals (including IL-4, IL-13, IL-10, and glucocorticoids) [7, 9, 10].

The dynamic balance of these MΦ phenotypes shifts during tissue damage and repair, with M1 dominant during early inflammatory and M2 during resolution and healing. Typical immune signalling pathways convert external stimuli into cytosolic events leading to resolution [11, 12], which can be achieved through materials providing biochemical cues or by therapeutic immunomodulatory molecules [13]. Many biomaterials have been shown to reduce harmful inflammation by modulating MΦs activity, both *in vitro, ex vivo* and *in vivo* [14–17]. For instance, engineered matrices enriched with immunomodulatory factors promoted enhanced bone formation by enhancing vascular invasion and early inflammation [15].

Hyaluronan (HA) is a linear, anionic polysaccharide that plays a key biological role in a range of human tissues [18] and has been widely used in biomaterials and tissue engineering [19–22]. Its effects depend on molecular weight (M_w_) and external environment, influencing biological pathways and paracrine signalling of immune cells through interactions with RHAMM and CD44 receptors [23, 24]. Notably, the CD44 receptor can distinguish HA M_w_, delivering signals that trigger different biological processes through clustering effects, as previously demonstrated *in vivo* and *in vitro* [25]. For example Rayahin *et al.* showed that high M_w_ (>1000 kDa) HA polarizes on murine MΦs (J774A.1) to an M2-like state, while low M_w_ HA (<1000 kDa) drives M1-like polarization [26]. Similarly, Lee *et al.* demonstrated that high M_w_ HA (>1250 kDa) inhibited pro-inflammatory responses in lipopolysaccharides (LPS)-stimulated RAW 264.7 murine MΦs, inducing anti-inflammatory effects in a concentration-dependent manner [27].

However, polarization in human-derived cells is more variable due to donor differences [28–30]. Contradictory findings on HA M_w_ effects have been attributed to different HA sources and endotoxin contamination [29, 31]. For example, high M_w_ HA (M_w_=1010–1800 kDa) has been shown to effectively polarize human monocytes (THP-1 cells) towards an M2-like state [32]. The overall polarization effects depend on cell type, source, HA concentration and M_w_ and endotoxin levels, although chemically functionalized HA versions, commonly used in tissue engineering, have not been as thoroughly investigated.

Tyramine-modified hyaluronan (THA) has emerged as promising biopolymer for therapeutic biomaterials, enabling rapid biofabrication through both enzymatic (horseradish peroxidase, H_2_O_2_) and light-based crosslinking (e.g. Eosin and green light irradiation) [33–35]. THA has also been used with collagen as an extrudable bioink to induce cellular alignment and migration [36]. Recently, our group showed THA (M_w_=280 kDa) hydrogels exhibited immunoinertness in neutrophil studies, showing no enhanced pro-inflammatory response compared to other materials [14].

Designing functional biomaterials that integrate immunomodulatory effects, biocompatibility and that allow for stable long-term polarization of MΦs is of considerable interest in tissue engineering and biofabrication fields [13]. Hence, in this study we evaluated the capacity of THA of two molecular weights: M_w_=280 kDa (LMW THA) and M_w_=1640 kDa (HMW THA) to polarize MΦs. THA was supplemented in cell culture medium at a concentration of 2 mg mL^-1^, which falls within the typical range observed in human knee synovial fluid [37]. This soluble signal is expected to: (i) mimic the release of THA from crosslinked hydrogels, where a 10% degradation rate of hydrogel containing 20 mg mL^-1^could release fragments at this concentration [38], or (ii) serve as a direct carbohydrate supplement in the medium. The polarization effects of the THA were tested on MΦs derived from two sources: genetically similar THP-1 cells and human peripheral blood mononuclear cells (PMBCs) from three donors. M1/ M2 polarization induced by THA supplementation were assessed using flow cytometry and multiplex ELISA for THP-1-derived MΦs while PBMCs-derived MΦs polarization was evaluated through immunofluorescent staining, gene expression analysis and ELISA.

## 2. Materials and methods

### 2.1. Synthesis and characterisation of tyramine-modified hyaluronan (THA)

THA of two molecular weights: M_w_=280 kDa (abbreviated throughout as LMW THA) and M_w_=1640 kDa (abbreviated throughout as HMW THA) were synthesized as previously described [34, 39]. Briefly, sodium hyaluronate (Contipro Biotechs.r.o., Czech Republic) was dissolved in deionized H_2_O at 1 w/v%. THA conjugates were prepared in a one-step reaction by addition of 1.25 mmol 4-(4,6-dimethoxy-1,3,5-triazin-2-yl)-4-methylmorpholinium chloride (DMTMM, TCI Europe N.V.) coupling agent and subsequently 1.25 mmol tyramine hydrochloride (Roth, 5234.3) was added dropwise to the solution. The reaction was carried out at 37 °C and under continuous stirring for 24 h. The product was purified via precipitation with 96% ethanol after the addition of 10 v/v% saturated sodium chloride. Several wash steps were performed, and the product kept under vacuum for 72 h, with first 24 h at 25°C and then for two final days at 40°C. For HMW THA, the purification and washing were repeated twice to ensure complete removal of trapped H_2_O and ethanol. Proton nuclear magnetic resonance spectroscopy (^1^H NMR, 300 MHz, Bruker Avance III) and UV absorbance measurements at 275 nm on Infinite® 200 Pro plate reader (Tecan, Männedorf, Switzerland) were performed to confirm the degree of substitution (DoS) of tyramine on HA (with DoS_LMW THA_= 6.01 ± 0.57 %, and DoS_HMW THA_= 5.99 ± 0.57 %). ^1^H NMR data is presented in ***Figure S1***. Loss of drying (LOD) was also evaluated. For that, 100 mg of THA was weighted into a glass vial and dried under vacuum at 105°C for 24 h. LOD was then calculated following the formula:

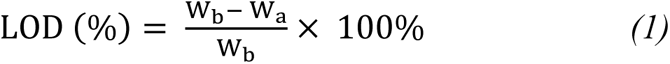

where W_b_ defines weight of THA before drying and W_a_ defines weight of THA after drying. The following values were obtained: LOD_LMW THA_= 9.8 ± 0.5 %, and LOD_HMW THA_= 12.8 ± 0.5%.

### 2.2. Endotoxin analysis

Bacterial endotoxins (lipopolysaccharides, LPS)-cell wall material from gram negative bacteria is a commonly known molecule that can activate immune cells [40]. To evaluate the LPS content of THA, the endotoxin content was measured using the Pierce™ Chromogenic Endotoxin Quant Kit (Thermo-Fisher) according to the manufacturer’s instructions. To evaluate THA, we adapted our previous protocol [14], and prepared enzymatically crosslinked THA hydrogels at 3.5 w/v% with 0.1 U mL^-1^ Horseradish peroxidase (HRP) 0.15 mM H_2_O_2_, the same for both LMW and HMW THA. The THA hydrogels were directly prepared in the wells and were incubated for 1 hour with endotoxin free H_2_O (Sigma-Aldrich). An endotoxin standard stock solution was prepared by adding endotoxin free H_2_O (Sigma-Aldrich) to the E. coli endotoxin standard vial to create a high standard (0.1-1.0 EU mL^-1^) and a low standard (0.01-0.1 EU mL^-1^). After 1 hour the supernatant H_2_O was harvested from the THA hydrogels and transferred to the pre-warmed readout plate at 37 °C as well as two standard curves were prepared on this plate. The amoebocyte lysate reagent was added to each well and the plates were incubated for 8 min at 37 °C. The reaction of the chromogenic substrate solution was stopped after 6 min at 37 °C by adding 25 v/v% acetic acid to each well and the optical density was measured at 405 nm using the Infinite® 200 Pro plate reader (Tecan, Männedorf, Switzerland). The obtained values were used to calculate the concentration based on the provided standard curve for the aforementioned concentration ranges, with R^2^=0.98 and R^2^=0.95 for low and high standards, respectively.

### 2.3. Quantification of the pro-inflammatory response of monocytes via NFKB expression

THP-1 monocytic cells with the NFκB-GFP reporter gene were kindly provided by Dr. Xin Xie (Core Technology Platform, New York University Abu Dhabi, UAE). The cells were cultured in RPMI-1640 media supplemented with 10% fetal bovine serum (FBS), 1% sodium pyruvate, 0.01% mercaptoethanol, and 1% penicillin/streptomycin at 37°C, 5% CO_2_, and 95% humidity. Cell culture media and supplements were purchased from Gibco™ (Thermo Fisher Scientific, Inc., Dreieich, Germany). To quantify the pro-inflammatory response, 5 × 10^5^ cells were cultured with or without at 2 mg mL^-1^ THA (10 times diluted from originally prepared THA hydrogels at 20 mg mL^-1^ with 0.1 U mL^-1^ Horseradish peroxidase (HRP) 0.15 mM H_2_O_2_), for 24 hours under standard cell culture conditions. As a positive control, the cells were treated with 10 ng mL^-1^ TNF-α (Biolegend, San Diego, CA, USA), known to activate NFκB signalling pathway. Cells were also treated with 10 ng mL^-1^ IFN-γ, as it is a known to confirm specificity of control because it does not trigger NFkB signalling pathway. Subsequently, the cells were analysed using an Attune NxT Flow Cytometer equipped with an autosampler (Thermo Fisher Scientific, Inc., Dreieich, Germany). Data analysis was performed using FlowJo software (Becton, Dickinson and Company, Franklin Lakes, NJ, USA). The geometric mean fluorescence intensity (gMFI) of the NFκB-GFP reporter was evaluated. Experiments were performed in 5 replicates.

### 2.4. MΦ differentiation from THP-1 cells and treatment with the soluble THA

For MΦ differentiation, 1 × 10^5^ THP-1 cells (ATCC, Manassas, VA, USA) were incubated with 300 nM phorbol 12-myristate 13-acetate (PMA; Sigma-Aldrich, Darmstadt, Germany) in RPMI-1640 media supplemented with 1% sodium pyruvate, 0.01% mercaptoethanol, and 1% penicillin/streptomycin but without fetal bovine serum (FBS), following previously published protocols [41, 42]. The cell culture media and supplements were acquired from Gibco™ (Thermo Fisher Scientific, Inc., Dreieich, Germany). After 6 h, the differentiation medium was removed, and the cells were allowed to rest for 24 h in RPMI-1640 without FBS and PMA. Subsequently, the differentiated MΦs were cultured either with or without 2 mg mL^-1^ THA (10 times diluted from originally prepared THA hydrogels at 20 mg mL^-1^ with 0.1 U mL^-1^ HRP 0.15 mM H_2_O_2_), for 48 h. As a control, MΦs were activated into pro-inflammatory (M1) MΦs for 48 h by adding 10 pg mL^-1^ lipopolysaccharide (LPS; Sigma, USA) and 20 ng mL^-1^ interferon-gamma (IFN-γ; Biolegend, San Diego, CA, USA). For anti-inflammatory activation, MΦs were induced into the M2a subtype with 20 ng mL^-1^ interleukin 4 (IL-4; Biolegend, USA) and 20 ng mL^-1^ interleukin 13 (IL-13; Biolegend, USA), or into the M2c subtype with 20 ng mL^-1^ interleukin 10 (IL-10; Biolegend, USA).

### 2.5. Quantitative analysis of THP-1-derived MΦs cell surface markers and their cytokine secretion profile

For analysis of MΦ cell surface markers, MΦs were detached from the cell culture plates using Macrophage Detachment Solution DXF (PromoCell, Heidelberg, Germany) following incubation with 2 mg mL^-1^ THA (10 times diluted from originally prepared THA hydrogels at 20 mg mL^-1^ with 0.1 U mL^-1^ HRP 0.15 mM H_2_O_2_). Before cell staining, they were incubated with Human TruStain FcX™ – Fc Receptor Blocking Solution (5 μL of blocking solution in 1000 μL of staining volume; Biolegend, San Diego, CA, USA) for 5 min under standard cell culture conditions. Subsequently, the cells were stained with mouse anti-human HLA-DR (clone: L243) conjugated with Brilliant Violet 421, CD163 (Clone: GHI/61) conjugated with PE-Cy7, mouse anti-human CD206 (Clone: 15-2) conjugated with Brilliant Violet 510, and DRAQ7 (a cell viability staining dye) for 30 min on ice. The antibodies and DRAQ7 dye were diluted in PBS at a 1:250 ratio. All antibodies used in this study were sourced from Biolegend, San Diego, CA, USA. The stained cells were then analysed using an Attune NxT Flow Cytometer equipped with an autosampler (Thermo Fisher Scientific, Inc., Dreieich, Germany), with compensation settings adjusted prior to analysis. Data analysis was conducted using FlowJo software (Becton, Dickinson and Company, Franklin Lakes, NJ, USA), focusing on the geometric mean fluorescence intensity of the cell surface markers. Experiments were performed in 5 replicates. For cytokine secretion profile analysis, cell culture supernatants were harvested following incubation with THA as already described. The cytokine levels were quantified using a bead-based multiplex immunoassay (LEGENDplex™ Human Essential Immune Response Panel; Biolegend, San Diego, CA, USA) designed for IL-4, IL-2, CXCL10 (IP-10), IL-1β, TNF-α, CCL2 (MCP-1), IL-17A, IL-6, IL-10, IFN-γ, IL-12p70, CXCL8 (IL-8), and TGF-β1, according to the manufacturer’s protocol. The samples were analysed on an Attune NxT Flow Cytometer (Thermo Fisher Scientific, Carlsbad, CA, USA). Data were processed using a 5-parameter curve fitting algorithm with LEGENDplex™ data analysis software (Biolegend, San Diego, CA, USA), and experiments were performed in 5 replicates.

### 2.6. MΦ differentiation from peripheral blood mononuclear cells (PBMCs) and treatment with the soluble THA

Primary human monocytes were isolated from buffy coats purchased from the Regional Blood Donation Service SRK Graubünden (Chur, Switzerland) from three human donors (D1, D2 and D3) using Ficoll density gradient separation. No donor information was received as per local regulations. Obtained peripheral blood mononuclear cells (PMBCs) were labelled with CD14 magnetic beads followed by magnetic-activated cell sorting (MACS) [43]. CD14^+^ monocytes cells were resuspended in RPMI medium supplemented with 10% FBS and then seeded onto tissue culture plastic at a density of 1.82 x 10^5^ cells well^-1^ for D1, and at a density of 3.0 x 10^5^ cells well^-1^ for donors D2 and D3, both in the presence of 20 ng mL^-1^ human macrophage colony-stimulating factor (hM-CSF) for 5 days. Medium (still containing hM-CSF) was exchanged on day 2. At day 5, hM-CSF-free medium was supplemented with 10 ng mL^-1^ TNF-α and IFN-γ for M1 MΦs and 10 ng mL^-1^ IL-4 for M2 MΦs for 24 h followed by a 24-hour rest in medium without cytokines (***Figure 3A***). M0-like MΦs control samples were cultured in medium without any cytokines (***Figure 3A***). Furthermore, on day 5, the same three polarization states were maintained (M0, M1, M2), either without any external influence of materials, or in the presence of the 2 mg mL^-1^ LMW THA and HMW THA supplemented in the cell culture medium (***Figure 3A***). Noteworthy, 2 mg mL^-1^ of both LMW and HMW THA remained liquid and did not result in a formation of any self-supporting physical hydrogels. On day 7, conditioned medium was collected, samples were washed two times with PBS and fixed with 4% formaldehyde for 20 min for immunofluorescent staining and stored in TRI-reagent for subsequent RNA isolation (***Figure 3A***).

### 2.7. Immunofluorescence analysis

D1 samples for immunofluorescent analysis were stained for nucleus with DAPI (NucBlue® Fixed Cell ReadyProbes® Reagent, ThermoFisher), and for phalloidin (ActinGreen™ 488 ReadyProbes® Reagent, ThermoFisher). Samples were firstly permeabilized with 0.25% Triton X-100 in PBS for 30 min and subsequently washed two times with wash buffer (0.05% Tween-20 in PBS). Samples were then firstly stained with phalloidin and left for 1 h in darkness. Subsequently, they were washed again two times with wash buffer, and finally incubated with DAPI for 10 min in darkness. Manufacturers’ instructions were followed for DAPI and ActinGreen dilutions (1 drop per 0.5 mL).

D3 samples for immunofluorescent analysis were stained for nucleus with DAPI (NucBlue® Fixed Cell ReadyProbes® Reagent, ThermoFisher), for phalloidin (ActinRed™ 555 ReadyProbes® Reagent, ThermoFisher), for MMR/CD206 (M2a MΦ surface marker) with Alexa Fluor® 700 (FAB25342N, R&DSystems, Bio-Techne), and for CD68/SR-D1 (a glycoprotein marker that is primarily found on the surface of MΦs) with Alexa Fluor® 488 (IC20401G, R&DSystems, Bio-Techne). Samples were firstly permeabilized with 0.25% Triton X-100 in PBS for 30 min and subsequently washed once with wash buffer. Samples were then firstly stained with phalloidin and left for 3 h in darkness. Subsequently, they were washed one time with wash buffer, and then incubated with blocking solution (1% BSA in 1x PBS) for 1 h. Then samples were washed one time with wash buffer and incubated with CD68 or CD206 staining (separate wells) pre-made at 8 µg mL^-1^ with 0.1% BSA in PBS overnight in darkness. Subsequently samples were washed one more time with wash buffer and incubated with DAPI for 10 min. Manufacturers’ instructions were followed for DAPI and ActinRed dilutions (1 drop per 0.5 mL). D2 samples were used to establish the appropriate staining protocol, subsequently used, and described above for D3 samples.

Samples were analysed using confocal microscope (Zeiss LSM800 Airyscan Confocal Microscope) with channels at 405 nm (DAPI), 561 nm (ActinRed), 515 nm (ActinGreen/CD68 Alexa Fluor 488) and 723 nm (CD206, Alexa Fluor 700).

Image analysis (nucleus cell area and circularity, cells area as well as CD68 area) was performed using ImageJ® on images taken using 10x objective. Only areas in the range from 5 to 120 µm^2^ and 90 to 400 µm^2^ were taken into consideration, for nucleus and cell, respectively. For CD68 analysis, 10 to 400 µm^2^ areas were taken into consideration. Smaller areas would be indicative of fragments whereas larger were excluded based on threshold found in manual analysis indicating more than 2 nuclei/cell of cells overlapping with each other/touching. The number of CD68^+^ and CD206^+^ was calculated by determining the total number of cells expressing CD68/CD206 and total number of DAPI per each obtained confocal image using Fiji (ImageJ2®). The average number of cells expression was then:

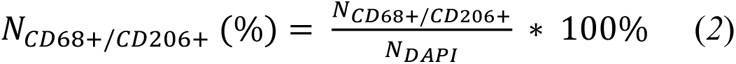

The employed method was semi-automative, with determining all staining in the area 10 to 400 µm^2^ in consideration and manually adding/subtracting the overlapping cells cases. The average number with presented standard deviation was calculated based on values obtained from 5-8 independent images taken for each group, as presented in Figure 4.

### 2.8. RNA isolation and RT-qPCR gene analysis of PBMCs-derived MΦs

To assess the polarization of MΦs, cells were collected and seeded in a 24-well plate at a concentration and with treatments described in section 2.6. RNA was isolated with salt precipitation. The obtained RNA content ranged from 420 to 1240 ng (***Table S1***). TaqManTM Reverse Transcription Kit was used to generate cDNA. Relative gene expression reactions were set up in 10 µL reaction mixes using TaqManTM MasterMix relevant human primers (***Table 1***), diethyl pyrocarbonate treated water (DEPC-water) and cDNA (10 ng). Real-time polymerase chain reaction (real-time PCR) was performed on two technical duplicates for n=4 (D1) or n=3 (D2 and D3) repeats, using QuantStudio 7 Flex. The relative gene expression was identified log_2_(fold change), where fold change = 2^-ΔΔCt^ value with RPLP0 as endogenous control and M0 MΦs with no supplemented material as control were used for normalization.

**Table 1.**
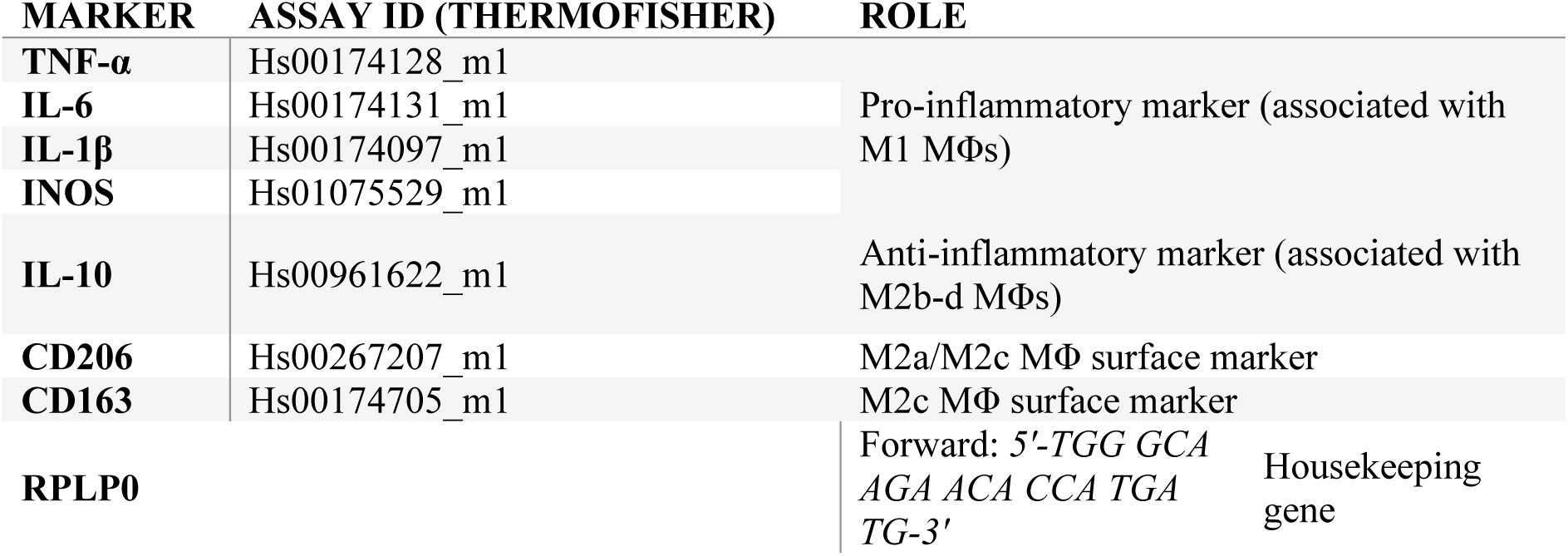
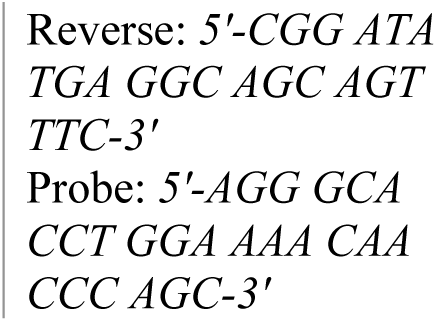
Human primer sequences for real-time PCR.

### 2.9. Enzyme-linked immunosorbent assay (ELISA) of PBMCs-derived MΦs

At the end of each polarization experiment, supernatant media was collected and kept for measurements of pro-inflammatory (IL-1β, IL-6, TNF-α), anti-inflammatory (IL-10) cytokines and anti-inflammatory surface receptor marker associated with M2c MΦs (CD163). The DuoSet ELISA systems were used according to the manufacturer’s instructions (DY201, DY206, DY210, DY1607, DY217B, R&D Systems, respectively). Absorbance values were measured on Infinite® 200 Pro plate reader (Tecan, Männedorf, Switzerland) at 450 nm with a background wavelength correction of 540 nm.

### 2.10. Statistical analysis

All results were analysed and presented with GraphPad Prism version 9.3.1. Results are shown as an average ± standard deviation (SD), with individual points marked. To perform statistical significance one-way analysis of variance (ANOVA) with Šídák’s multiple comparisons, Tukey’s multiple comparisons test, or Kruskal-Wallis multiple comparisons or two-way analysis of variance (Anova) with mixed effects were used and noted in each case in the Figure description. A statistically significant results were considered as of p<0.05 (ns ->0.05, * - <0.05, ** - <0.01, *** - <0.005 and **** - <0.001).

## 3. Results and discussion

### 3.1. Endotoxins analysis

To eliminate the possible effects stemming from the endotoxins, we confirmed that their measured levels were 0.27 ± 0.07 U mL^-1^ and 0.26 ± 0.04 U mL^-1^ for LMW THA and HMW THA, respectively, below the limit for clinical medical devices of 0.5 EU mL^-1^ set by the USA Food and Drug Administration (FDA) [44]. Given that endotoxin levels were similar for both LMW and HMW THA, we anticipate that any unravelled differences due to THA in polarisation experiments discussed throughout this article will stem solely from the difference in M_w_.

### 3.2. Immunological effects exerted on genetically similar monocytic THP-1 cells-derived MΦs

Firstly, we evaluated the effects of supplemented THA on genetically similar THP-1 monocytic cells that carried the NFκB-GFP reporter gene and we tested whether supplementation has any capacity to activate NFκΒ signalling pathway (***Figure S2***), which is well correlated with the induction of pro-inflammatory responses [45]. Genetically similar THP-1 cells enable screening of immunomodulation potential by minimizing any biological variability stemming from primary human cells. Cells treated with IFN-γ, which is known not to trigger this pathway, as well as cells supplemented with LMW and HMW THA did not show any significant upregulation as compared to untreated cells, and hence do not trigger any obvious pro-inflammatory response in this model under the tested conditions. Out of all tested samples, only when cells were treated with TNF-α, this pathway was activated as evidenced by significantly higher geometric mean fluorescent signal intensities (p<0.005) as compared to untreated cells (***Figure S2***) [46].

#### 3.2.1. Flow cytometry profiling of THP-1 derived MΦ phenotypes

Next, following the protocol established by Sapudom *et al.* [41], we investigated the general polarization capacity of THA on THP-1-derived MΦs (***Figure 1A***). MΦ phenotypes were characterized by flow cytometry using four specific surface markers: HLA-DR for pro-inflammatory MΦs (M1), CD105 for anti-inflammatory MΦs (M2), and subsequently CD163 for anti-inflammatory MΦs (M2c) and CD206 for anti-inflammatory MΦs (primarily for M2a/M2c) sub-phenotypes [47, 48]. Control M1 and M2 subtypes (M2a and M2c) are also presented in all graphs as horizontal lines. We observed significantly higher abundance of HLA-DR surface markers (p<0.01) for cells encapsulated with LMW THA (***Figure 2B***). Pro-inflammatory MΦs typically exhibit HLA-DR expression levels approximately 5-to 10-fold higher than those seen in anti-inflammatory MΦs, whether derived from PBMCs or THP-1 cells [48, 49]. Hence, it appears that LMW THA stimulates a pro-inflammatory polarization capacity in our model. On the other hand, we did not see any significant difference for HMW THA as compared to M0 control (***Figure 2B***). We then noticed significantly higher abundance of CD105^+^ cells (p<0.01) when encapsulated both with LMW and HMW THA (***Figure 1C***), generally indicating cell involvement in angiogenesis and tissue repair processes [42, 47].

**Figure 1.**
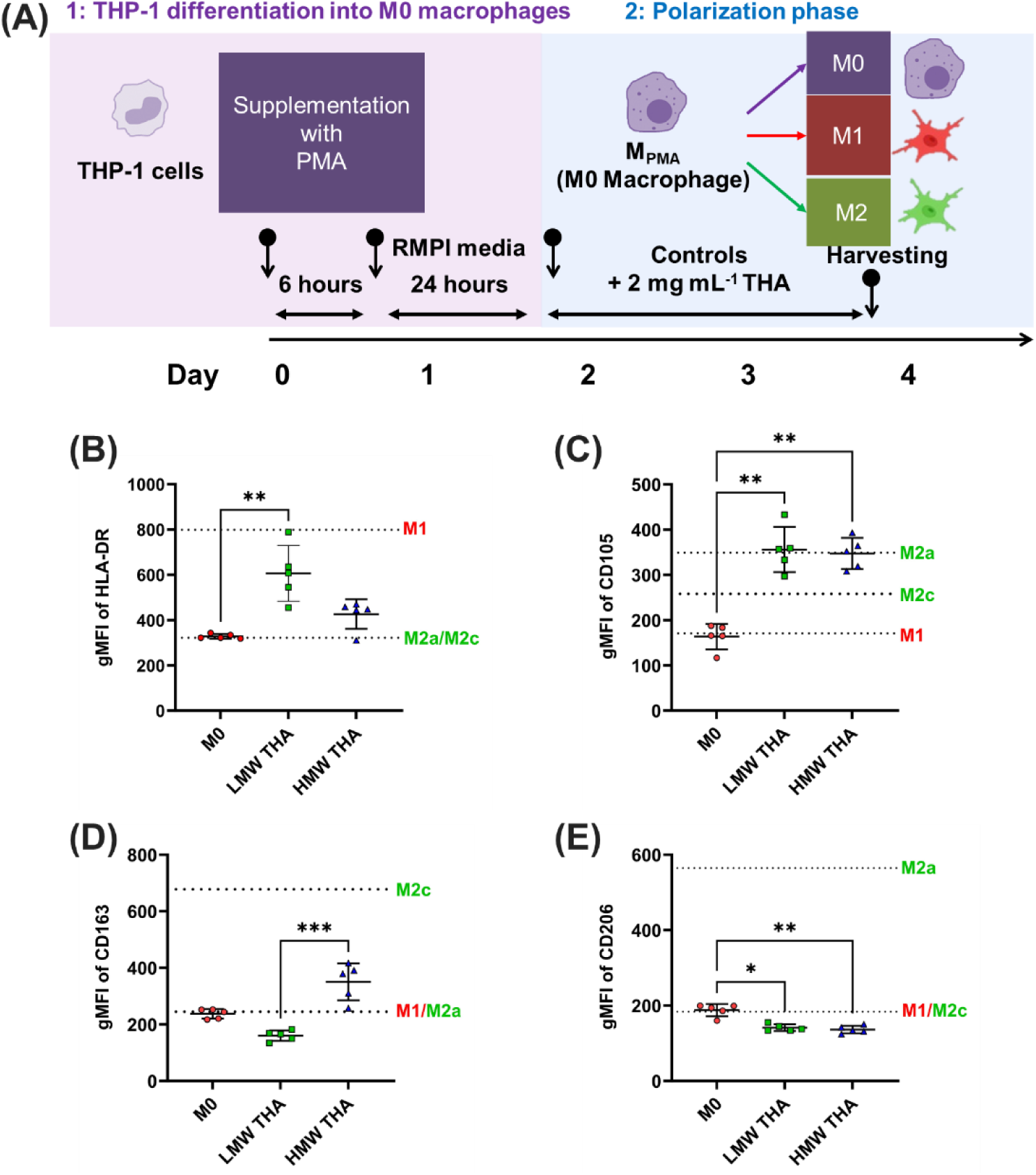
**(A)** Scheme summarizing the differentiation protocol of THP-1-derived MΦs. THP-1 cells were differentiated into M0 MΦs by treatment with 300 nM phorbol-12-myristate-13-acetate (PMA) for 6 hours, followed by resting for 24 h in cell culture media. Subsequently, M0 MΦs were cultured in the presence of the 2 mg mL^-1^ supplemented in the cell culture medium for 48 h. Quantitative analysis of cell surface markers of **(B)** HLA-DR, **(C)** CD105, **(D)** CD163 and **(E)** CD206. Geometric mean fluorescence intensity (gMFI) was plotted. Black line and error bar in the plot represent mean and standard deviation, respectively. In all cases **(B-E)**, horizontal dashed lines depict average values for the control polarized MΦs obtained as indicated in methods section. Statistical analysis in **(B-E)** was done by Kruskal-Wallis multiple comparisons test. A statistically significant results were considered for p<0.05 (* -<0.05, ** - <0.01, *** - <0.005 and **** - <0.001).

**Figure 2.**
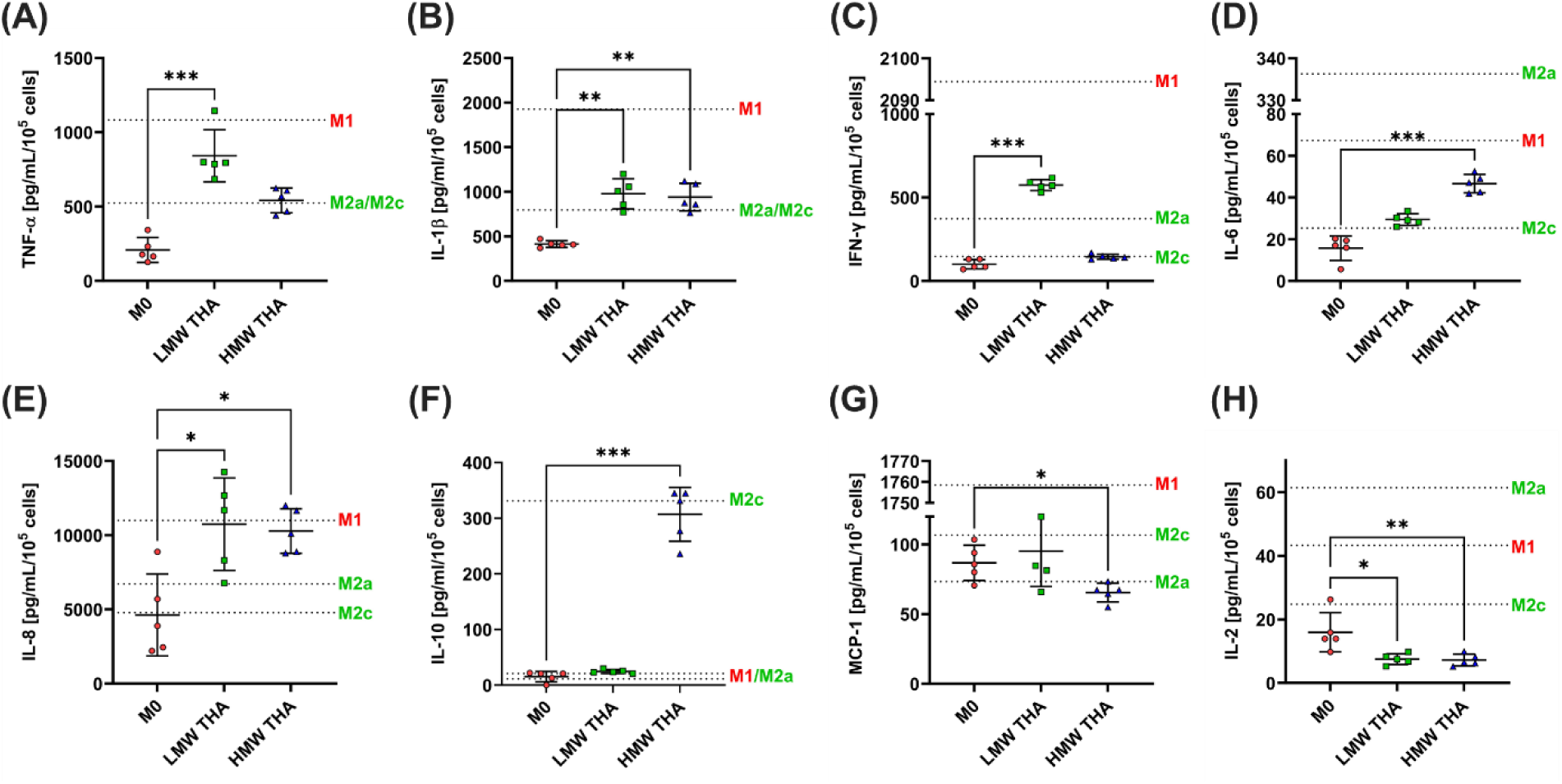
Cytokine secretion profile using multiplex bead-based ELISA on THP-1 derived MΦs. Significant changes were found only in 8 cytokines, presented here: **(A)** TNF-α, **(B)** IL-1β, **(C)** IFN-γ, **(D)** IL-6, **(E)** IL-8, **(F)** IL-10, **(G)** MCP-1, **(H)** IL-2. In all cases **(A), (B)**, and, horizontal dashed lines depict average values for the control polarized MΦs obtained as indicated in methods section. Statistical analysis was done by Kruskal-Wallis multiple comparisons test. A statistically significant results were considered for p<0.05 (* -<0.05, ** - <0.01, *** - <0.005 and **** - <0.001).

Given the importance of the M2 subtypes in tissue resolution, next we evaluated specific surface receptors, including CD163 (***Figure 1D***) and CD206 (***Figure 1E***), known to be associated primarily with M2c and M2a subtypes, respectively. Significant upregulation of CD163^+^ receptor (p<0.005) was found for cells supplemented with HMW THA, as compared to LMW THA (***Figure 1D***). At the same time, significantly lower abundance of CD206^+^ cells were observed for both LMW THA (p<0.05) and HMW (p<0.01), as compared to untreated cells (***Figure 1E***).

In summary, flow cytometry suggests that both LMW and HMW THA treatments lead to upregulation of CD105, suggesting a common involvement in tissue repair and angiogenesis, possibly stemming from the common chemistry of both molecules. However, they differ in their effects on HLA-DR, CD163, and CD206 expression, indicating differences in pro-inflammatory activation (M1) and M2c MΦ polarization, both deviating from standard M2a MΦ subtype. This points towards the fact that HMW THA may stimulate differentially the THP-1 derived MΦs towards a number of states, including the tissue reparative state and promotion of the healthy restoration of normal tissue architecture [42].

#### 3.2.2. Multiplex ELISA profiling of THP-1 derived MΦ phenotypes

In the next step, we evaluated the secretion profiles of 13 most common cytokines released by THP-1 derived MΦs using multiplex bead-based ELISA: IL-2, IL-4, IP-10 (CXCL10), IL-1β, TNF-α, MCP-1 (CCL2), IL-17A, IL-6, IL-10, IFN-γ, IL-12p70, IL-8 (CXCL8), and TGF-β1. Significant changes were found in eight of the tested cytokines:

- pro-inflammatory - TNF-α (***Figure 2A***), IL-1β (***Figure 2B***), *IFN-γ* (***Figure 2C***), IL-6 (***Figure 2D***), IL-8 (***Figure 2E***),
- anti-inflammatory - IL-10 (***Figure 2F***),
- regulatory – MCP-1(***Figure 2G***) and IL-2 (***Figure 2H***).

Higher levels of TNF-α (***Figure 2A***), IL-1β (***Figure 2B***), IFN-γ (***Figure 2C***), IL-8 (***Figure 2E***), and MCP-1 (***Figure 2G***) suggest that LMW-THA supplementation generated MΦs with pro-inflammatory phenotype and M1-like characteristics. These cytokines are typically associated with pro-inflammatory responses and activation of M1 MΦs. On the other hand, lower levels of IL-2 (***Figure 2H***) suggest a deviation from the typical M1 phenotype, as IL-2 is generally associated with T-cell activation and is not typically produced by M1 MΦs. Nevertheless, overall, the obtained cytokine profile suggests an M1-like pro-inflammatory phenotype for the MΦs supplemented with LMW THA and aligns with flow cytometry data, pointing towards partly deviated M1-like phenotype.

On one hand, similarly to LMW THA supplementation, HMW THA showed higher levels of IL-1β (***Figure 2B***) and IL-8 (***Figure 2E***), suggesting polarization towards a pro-inflammatory phenotype with M1-like characteristics. On the other hand, higher levels of IL-6 (***Figure 2D***) and IL-10 (***Figure 2F***) both suggest a deviation towards an M2-like phenotype with anti-inflammatory and immunoregulatory functions. IL-6 can be produced by both M1 and M2 MΦs, while IL-10 is typically produced by M2b, M2c or M2d subtypes and it is associated with anti-inflammatory responses key for the remodelling of external matrix [47, 50]. This goes in line with lower levels of MCP-1(***Figure 2G***), which suggest a potential decrease in chemotactic activity and recruitment of monocytes/MΦs, which indicates a shift away from the pro-inflammatory M1 phenotype. Lower levels of IL-2 (***Figure 2H***) also suggest similarly to the LMW THA supplementation a potential deviation from the typical M1 phenotype. Overall, the cytokine profile for HMW THA supplementation suggests a mixed phenotype with characteristics of both M1 and M2 MΦs, possibly indicating a transitional or heterogeneous population of MΦs.

Wang and Bratlie provided guidelines for MΦ polarization as a function of used polymer chemistry properties [51]. They argued that chemistry leading to more H-bonding capacity and higher hydrophilicity stimulates MΦs towards an M1 state, whereas overall charge plays a role in M2 modulation. Given our observations we currently cannot decouple the exact role of the tyrosine-chemical modification on the obtained profiling and point out that to some extent it may be influencing the outcome. Considering data from both flow cytometry and Multiplex ELISAs, the LMW THA supplementation induces a more pro-inflammatory M1-like phenotype in the THP-1-derived MΦs, while the HMW THA supplementation leads to a more intricate profile with some features of both M1 and M2 phenotypes, suggesting either a heterogeneous population or a transitional state. The observed profiles for LMW and HMW THA are however generally consistent with what might be expected for the treatment with non-modified low molecular weight hyaluronic acid (LMW-HA) and high molecular weight hyaluronic acid (HMW-HA), respectively [23, 52], indicating that the tyramine modification with low degree of substitution does not significantly alter HA’s biological response.

### 3.3. Immunological effects exerted on human donor-derived PMBCs-derived MΦs

#### 3.3.1. Cellular architecture parameters analysis

We then aimed to investigate the polarisation obtained with supplementation of THA molecules on primary peripheral blood mononuclear cells (PBMCs)-derived MΦs (***Figure 3A***). Primary human-derived cells are generally recognized as more clinically relevant because they better preserve tissue characteristics [53]. This importance has also been noted for MΦs, albeit with their large heterogeneity [54]. First, at day 6 of our protocol (***Figure 3A***) after the cytokine’s depletion, we evaluated using bright field microscopy the preliminary changes in cellular shape as a function of all polarization states and supplemented THA (***Figure S3***). Indeed, qualitatively more elongated cells were obtained across biochemically induced M1 and M2 states, indicative of cellular changes. Moreover, more cellular protrusions (i.e. filopodia) were observed across all M1 groups, in line with previous observations for M1 type of MΦs [55]. To confirm these findings, we firstly stained F-actin filaments and DAPI and performed immunofluorescence assay (on donor D1, from samples harvested from day 7 of polarization protocol) with semi-quantitative image analysis (***Figure 3B-E***). Fluorescent images confirmed similar findings that MΦs polarised biochemically *in vitro* to an M1 phenotype exhibited distinctly more lamellar processes, were of more elongated shape, whereas an M2 phenotype resulted in cells that were partly similar in shape to unpolarised M0 MΦs, with fewer lamellar processes. Furthermore, MΦs cultured in the presence of LMW THA appeared to have more filopodia and processes stemming from cellular bodies compared to control and HMW THA groups (***Figure 3B***). This trend has been confirmed with the significant differences in cellular areas between M0 and M1, and M1 and M2 groups (p<0.001, ***Figure 3C***), indicating a possible influence of LMW THA on the polarization state of obtained MΦs. A non-statistically significant trend for highest cellular area was observed for M1 phenotype, concurring with some previous observations [56]. For comparison, there were no significant differences noted for cells in either control or HMW THA groups across all polarization states (***Figure 3C***). For control group, there were also no statistical differences in area of nucleus between all polarization states (***Figure 3D***).

**Figure 3.**
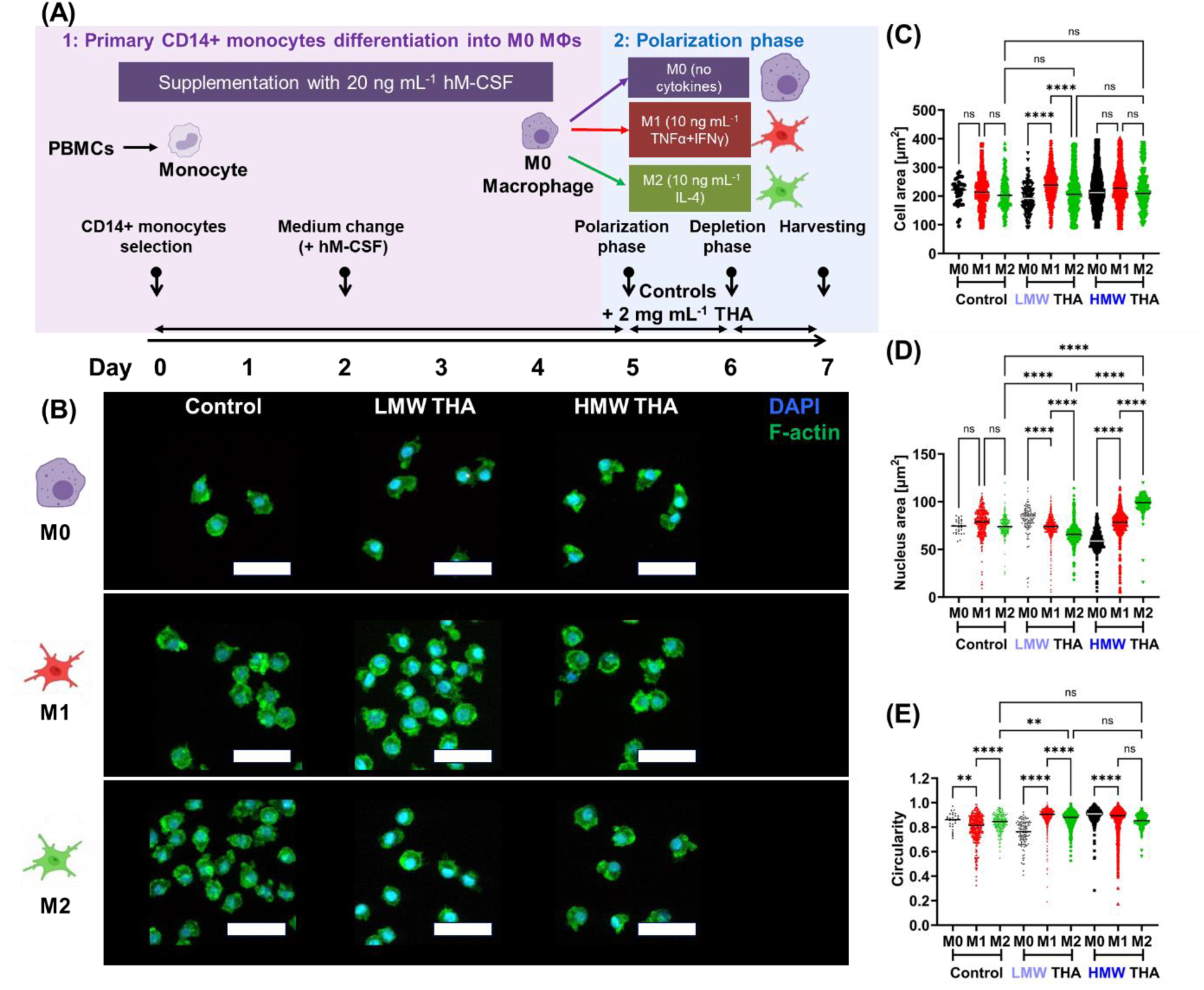
(A) Differentiation protocol and subsequent characterisation of peripheral blood mononuclear cells (PBMCs)-derived MΦs. PBMCs were differentiated into M0 MΦs over 5 days period by supplementation with 20 ng mL^-1^ human macrophage colony-stimulating factor (hM-CSF). At day 5, differentiated M0 MΦs were polarized by medium supplemented with 10 ng mL^-1^ TNF-α and IFN-γ towards an M1 state and 10 ng mL^-1^ IL-4 towards an M2 state for 24 h, followed by a 24-hour rest in medium only, without supplementation. M0-like MΦs control samples were cultured in medium without any cytokines. The same three polarization states were maintained (M0, M1, M2), either without any addition (control), or in the presence of the 2 mg mL^-1^ LMW THA and HMW THA supplemented in the cell culture medium at day 5. **(B-E) Immunofluorescence characterisation of peripheral blood mononuclear cells (PBMCs)-derived MΦs. (B)** Immunofluorescence staining of nucleus by DAPI (**blue**) and F-actin by Phalloidin (**green**) for all tested groups. Images were digitally enlarged to higher magnification to enable visualization of individual cells. Original images with lower magnification are provided in supporting information **Figure S3**. Each raw represents different polarization state (M0, M1 and M2) and columns represent a control and samples supplemented with 2mg mL^-1^ LMW or HMW THA. Scale bar = 50 µm. ImageJ analysis of obtained for at least 100 cells for all groups of: **(C)** cell area, **(D)** nucleus area and **(E)** nucleus circularity. Analysis by one-way ANOVA with Šídák’s multiple comparisons test. A statistically significant results were considered for p<0.05 (* - <0.05, ** - <0.01, *** - <0.005 and **** - <0.001).

Currently there exists a small number of studies directly comparing nucleus size among M0, M1, and M2 macrophage subtypes [55]. This report [55] suggests a possible decrease of nuclear area for M1 phenotypes, although no statistical significance was provided. Overall, while nucleus size variations between M0, M1, and M2 MΦ subtypes have not been extensively studied, it is plausible that differences in activation state and functional phenotype may influence nuclear characteristics. Some recent studies [57] have utilised artificial intelligence (AI) and image processing capabilities on large datasets of microscopical images from different MΦ polarization states. The authors suggested several descriptors (area, length, width, circularity, aspect ratio, roundness and solidity) of MΦ cellular morphology and indeed found significant differences noted between states. In our model, supplementation of LMW and HMW had a measurable influence, with nuclei area decreasing from M0 to biochemically induced state M1 and to M2 (p<0.001) in LMW THA group, and the nuclei area increasing from M0 to biochemically induced state of M1 and to M2 in the presence of HMW THA (***Figure 3D***), showing opposing trends for LMW and HMW THA groups.

The AI-based study of multiple MΦ subtypes [57] has indicated that total cell circularity may be one of the key varied cellular factors for the M2a, but without significant differences noted between M1 and M0 groups; however, circularity of nucleus across different MΦ subtypes was not evaluated. In our case in the control group, the nucleus circularity of M1 MΦs was found lowest (***Figure 3E***, p<0.01), which concurred with previous observations of human primary MΦs on fibrous architectures [58]. This trend of nucleus circularity was modulated by the presence of LMW THA, with M0 group having lower nucleus circularity, similar to that of M1 in control group, indicating potential pro-inflammatory influence of LMW THA (***Figure 3E***). However, M1 group in LMW THA retained higher nucleus circularity, similar to M0 in control group. This indicates that the effects stemming from simultaneous supplementation of LMW THA and 10 ng mL^-1^ TNF-α and IFN-γ of polarization cytokines might be mitigating each other. Although this analysis provides useful insights into modulatory potential of M_w_ of THA, we acknowledge its effects may be limited and, in this case, donor (D1) specific. In future, we see approaches similar to [57] undertaken on much larger sets of obtained images and across multiple donors in a machine-learning approach in anticipation of elucidation of donor-specific features. This work is however out of the scope of the current manuscript.

#### 3.3.2. CD68 and CD206 surface receptors analysis

To further confirm full differentiation of monocytes into MΦs and investigate the propensity for M2 polarisation in PMBCs-derived MΦs, we performed immunofluorescent staining using CD68 [10, 59] and CD206 as M2 marker (***Figure 4***) [47]. All groups had nearly 100% expression of CD68^+^ with no significant differences detected between any groups (***Figure 4H***), proving successful differentiation into MΦs utilizing our protocol. CD68 is typically located in the endosomal/lysosomal compartment however it has been shown to rapidly shuttle to the cell surface, particularly for M1 MΦ phenotype [10, 59]. We then calculated the area of staining per cell to provide more quantitative overview of the possible localization of the staining, with larger values possibly indicating extension beyond the cytoskeleton, whereas lower values possibly indicating retainment within endosomal compartment. As expected, the largest CD68 staining area was observed in M1 polarization group in control cells (***Figure 4G***). Across all polarization groups supplemented with LMW THA, CD68 was higher than for M0 control group, with highest CD68 area per cell noted for M2 group (***Figure 4G***). In M0 and M1 states supplemented with HMW THA we observed the highest area of CD68 expression per cell (p<0.001 as compared to M0 control), whereas in M2 HMW THA case, lowest area (***Figure 4E*** and ***4G***). M0 group supplemented with HMW THA also showed significantly higher area of CD68 staining than M2 group supplemented with LMW THA (p<0.001, ***Figure 4G***). However, this effect was significantly supressed by synergetic action with supplemented 10 ng mL^-1^ IL-4, in which case, a potential retention of CD68 in the endosomal compartment could be possible, with lowest area per cell observed across all tested groups (***Figure 4G***).

**Figure 4.**
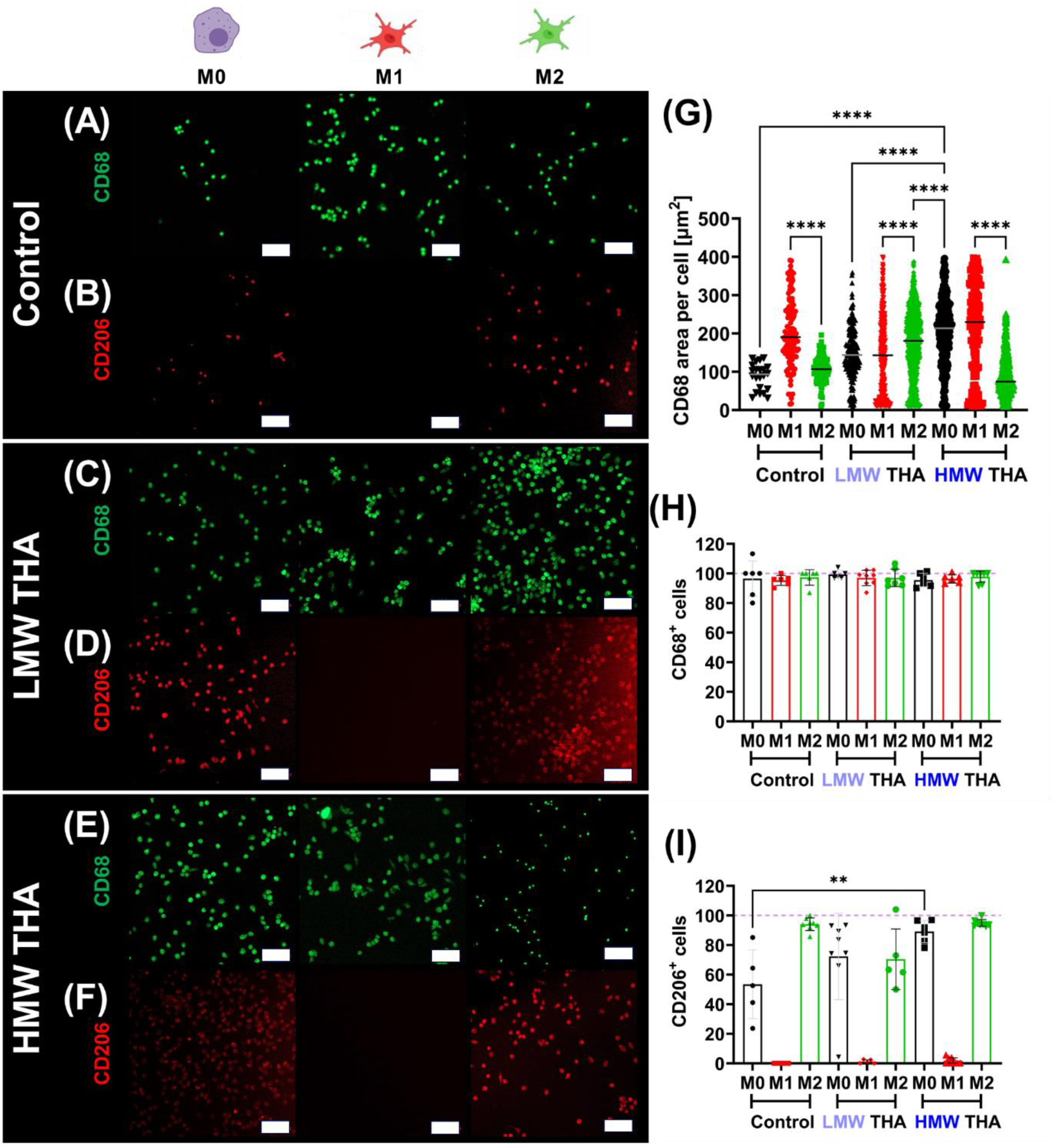
Surface marker characterisation of D3 peripheral blood mononuclear cells (PBMCs)-derived MΦs. **Immunofluorescence staining** of CD68 (**green**) is shown in panels **(A)**, **(C)**, and **(E)**; whereas staining of CD206 (**red**) is shown in panels **(B)**, **(D)**, and **(F)**. Each column represents different polarization state (M0, M1 and M2) as indicated on the top. **(A)** and **(B)** represent panels for a control, **(C)** and **(D)** represent samples supplemented with 2mg mL^-1^ LMW THA, and **(E)** and **(F)** represent samples supplemented with 2mg mL^-1^ HMW THA. Scale bars for all images = 100 µm. All images containing full images including staining of nucleus by DAPI (**blue**) and F-actin by Phalloidin (**orange**) are presented in **Figure S5**. **(G)** ImageJ analysis of obtained images for all groups of CD68 area per cell. Representative images are presented in **Figure 4A**, **C**, **E**, in main manuscript. Number of **(H)** CD68^+^ and **(I)** CD206^+^ positive cells calculated from the obtained images. Analysis by one-way ANOVA with Šídák’s multiple comparisons test. A statistically significant results were considered for p<0.05 (* - <0.05, ** - <0.01, *** - <0.005 and **** - <0.001).

Our data on THP-1-derived MΦs suggests that CD206 is a downregulated receptor (***Figure 1E***) when supplemented with either LMW or HMW THA. Noting that differential regulation of CD206 expression between THP-1-derived MΦs and PBMCs-derived MΦs in response to a given supplementation could stem from a combination of differences in cellular origin, underwent mutations to achieve immortalization, genetic/epigenetic factors, as well as microenvironmental cues, we also calculated numbers of CD206^+^ cells for PBMC-derived MΦs (***Figure 4I***). Expression of CD206 was reached nearly 100% in M2 polarization state in control group (94.0 ± 4.2 %) and M2 group supplemented with HMW THA (94.9 ± 2.4 %), (***Figure 4B, 4F, 4I***). Lower numbers of CD206^+^ cells (70.4 ± 20.5 %) were obtained for M2 polarization state supplemented with LMW THA (***Figure 4D*** and ***4I***), indicating the LMW THA may decrease of M2 MΦs population in biochemically induced M2 polarization state. All M1 states did not show any expression of CD206 (0-2%), as expected for all the three tested groups (control, LMW and HMW THA, ***Figures 4B, 4D, 4F***, and ***4I***).

The M-CSF differentiation factor, used in our study for MΦ maturation, has been reported to support a more M2-like phenotype [60, 61]. Indeed, we observed some % of M0 control group cells expressing CD206 (53.41± 23.2 %, ***Figure 4I***), pointing to the possibility (and in our case a limitation) of MΦs existing on a heterogenous spectrum across M0 and M2 states. The use of alternative GM-CSF differentiation factor is also known to induce the heterogeneous population of M0/M1 MΦs [60, 61], thereby too introducing a bias and being a limiting factor. Nevertheless, from our confocal microscopy, we detected significantly higher number of CD206^+^ cells when supplemented with HMW THA (p<0.01, 89.1 ± 7.6 %), but not significantly different with LMW THA (72.3 ± 29.2 %) (***Figure 4I***). This indicates that HMW THA supplementation increases the number of M2 states in PBMCs-derived MΦs, whereas LMW THA suppressed number of biochemically induced M2 states and retains heterogenous populations similar to M0 control group. Hence, our immunocytochemistry results only point to the somewhat more complex M0/M1/M2 heterogenous roles of supplemented LMW THA and HMW THA compared to the control groups. One limitation of our study is that the analysis was carried out on one single donor (D3), with outlook towards envisaged AI/machine learning methods on a much larger dataset of images in future. To unravel more complex dependence on multiple donors, we next run RT-qPCR gene analysis of some of the common markers, including CD206 gene expression.

#### 3.3.3. Gene expression analyses – intra versus inter-donor variability

First, we evaluated a panel of markers (***Table 1***) consisting of pro-inflammatory markers (TNF-α, IL-6, IL-1β, iNOS), an anti-inflammatory marker (IL-10) and M2 surface markers (CD206 and CD163), on donor D1 samples (***Figure S6*** and ***Figure S7A, D, G***). Notably, although not significantly different, all M1 groups were characterized by the upregulation of TNF-α (***Figure S7A***). Out of all pro-inflammatory markers we only detected significant differences in TNF-α between all M1 groups (control, supplemented with both LMW and HMW THA) and M0 supplemented with HMW THA (***Figure S7A***), with no significant differences observed between any of the groups for the remaining pro-inflammatory markers (***Figure S6***). The M0 group supplemented with HMW THA had lowest values across all groups, suggesting the possible influence in its reduction of inflammation, as already showed by our investigations using THP-1 derived MΦs. We also did not detect any significant differences between any groups in anti-inflammatory IL-10 marker (***Figure S7D***), with only miniscule trends observed. We obtained slightly lower values for the M1 groups and slightly higher values for the M2 groups, in general in line with the expectations for the classical polarization. Finally, we saw clear trends in gene expression of CD206 surface markers, with downregulation of all M1 groups and upregulation of all M2 groups (***Figure S7G***). The only significantly detected differences were for M2 group supplemented with LMW THA, which appeared to have much higher expression compared to all M1 groups (***Figure S7G***). However, we do note the lack of statistical difference between any M2 groups.

Given the emerging picture of gene expression in D1 (which we run in first instance, ***Figure S6*** and ***Figure S7A, D, G***), to fully evaluate intra-donor variability, we then run full gene expression analysis on these four key markers: TNF-α, IL-10, CD206 and CD163, with the upregulated TNF-α predominantly representing M1 state, upregulated IL-10 and CD206 representing M2a and upregulated CD163 M2c state [62] on all three donors (***Figure 5*** and ***Figure S7***). The full individual panel of all markers is provided in ***Figure S7***, whereas here we discuss general trends across all donors to confirm the overall effects of supplementation of THA molecules. Even though we observed upregulation in TNF-α, for LMW THA and downregulation for HMW THA, these were found not statistically significant across all donors (***Figure 5A***) as well as in mixed effects analysis. We could note slight tendencies of upregulation in M0 groups supplemented with LMW THA in all donors (not significant) (***Figure 5A***) concurring with the data obtained for THP-1-derived MΦs, but also possibly indicating weaker effects in PBMCs-derived MΦs. It appears that D1 seemed to differ from the other two donors. In D2 and D3 HMW THA supplementation had similar expression to M0 state (lack of significance, ***Figure 5A***). We also note that observed variations in regulation of TNF-α across donors may come from both environmental and genetic factors, including gender, white blood cells, tobacco consumption, and cholesterol levels [63]. These factors are not provided or controlled in the received blood donations, so drawing any decisive conclusions remains limited.

**Figure 5.**
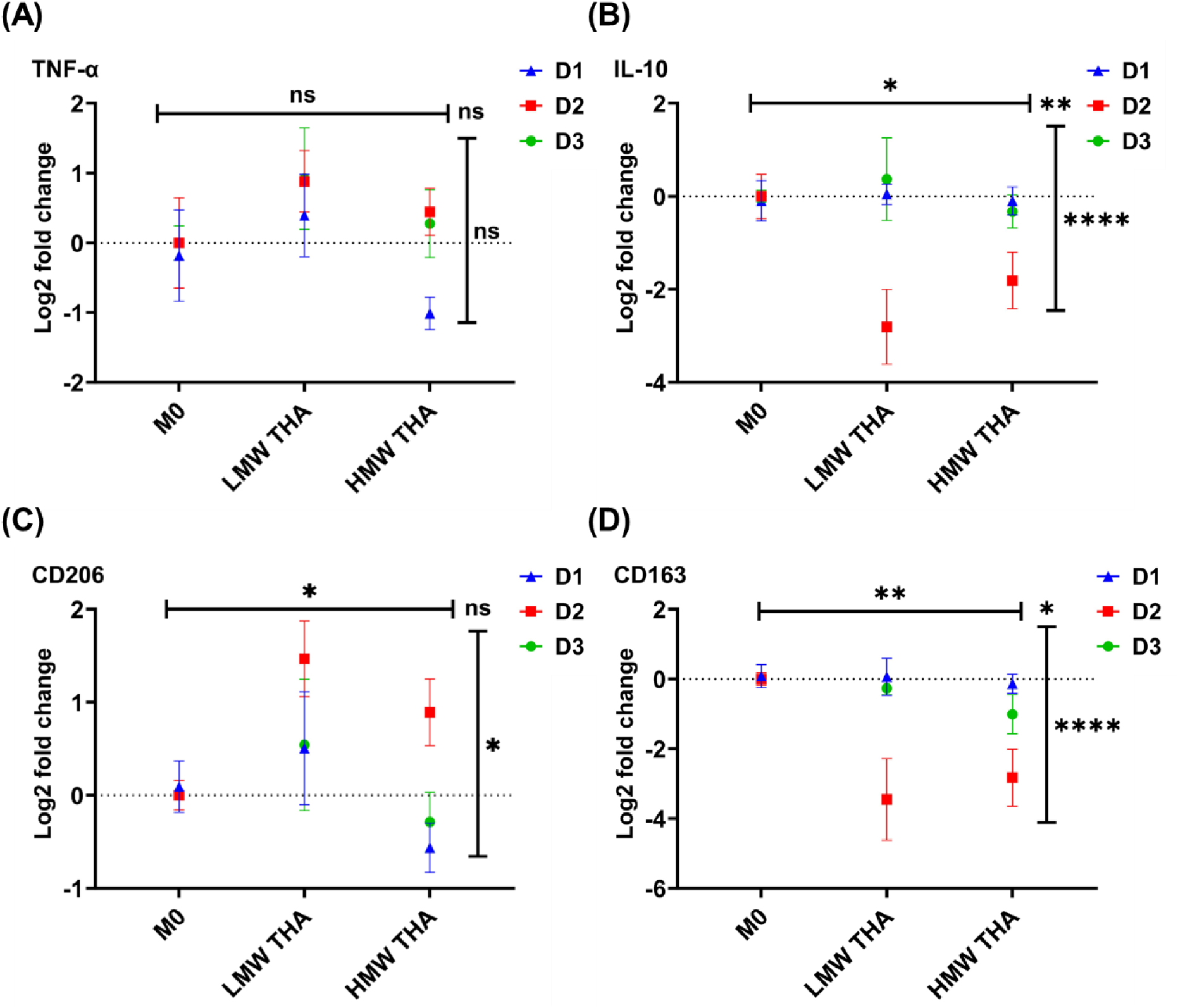
Log2 fold change gene expression of **(A)** TNF-α, **(B)** IL-10, **(C)** MRC1 (CD206), and **(D)** CD163, measured by RT-qPCR for cells without any material conditioning (control) or encapsulated with 2 mg mL^-1^ LMW or HMW THA. Data represent average ± SD value from n=4 for D1 and n=3 for D2 and D3 from individual experiments. Gene expression is represented as fold change compared to RPLP0 (a housekeeping gene) of the M0 control group. Mixed effects model analysis with Geisser-Greenhouse correction was performed. A statistically significant results were considered for p<0.05 (* - <0.05, ** - <0.01, *** - <0.005 and **** - <0.001), with horizontal dash representing significance across supplementation/material, vertical across donors and corner value provides significance of their interaction.

Unlike in the case of THP-1-derived MΦs where we measured high amount of released IL-10 from samples supplemented with HMW THA, for PBMCs-derived MΦs we did not observe any significant upregulation of IL-10 gene (***Figure 5B***). In fact, we observed lower expression of IL-10 for one donor (D2, ***Figure 5B***) when supplemented with both LMW and HMW THA. We also noticed some donor susceptibility to the typical biochemical induction of M1 and M2 states in the presence of LMW and HMW THA (***Figure S7E-F***). This, if anything, suggests a possible deviation from other M2 subtypes than M2a [47, 50]. One possibility is that THA of both M_w_ (***Figure 5E-F***, ***Figure S7D-F***) influenced the typical JAK/STAT IL-10 production metabolic pathways in PBMCs-derived MΦs, and that certain donors may have propensity for this pathway’s dysregulation [64] (***Figure 5B***, ***Figure S7D-F***). The mixed effects statistics indicated significance across supplementation, donors and their interaction (***Figure 5B***). The interpretation of the overall IL-10 gene regulation data for PBMC-derived MΦs is complex as human IL-10 regulation has been shown to be affected by interindividual variation in gene expression [65, 66]. We therefore believe that responses that we see in our data (p<0.001) stem directly from the variation introduced by host genetic factors, which are not pre-screened in received blood donations.

In the case of CD206 surface marker, we noted significance across supplementation and across donors, but not in their interaction (***Figure 5C***). For biochemically induced states, the CD206 upregulation was evident and coherent across all donors, with all M2 groups upregulated as compared to M1 groups (***Figure S7G-I***). By comparison of groups supplemented with LMW and HMW THA, in general LMW THA induced a slightly higher upregulation, whereas HMW THA variable regulation, with upregulation in D1 and downregulation in donors D2 and D3 (***Figure 5C***). In all cases differences are minimal. Although CD206 is typically associated with M2a/M2c states of MΦs, it has been shown that its activation may also be associated with response to the overall inflammatory state [67], which may be the case of LMW THA noted here for some donors. The slight increase of CD206^+^ cells as compared to M0 control has been shown in the immunofluorescence staining (***Figure 4I***). The high number of CD206^+^ cells in HMW THA group noted in immunofluorescence (***Figure 4I***) compared to gene expression of this marker (***Figure 5C***) is indicative of the different time scales for gene expression and protein synthesis, and of the different information obtainable at the mRNA and protein level. To deconvolute temporal effects of the THA supplementation, further studies outside of the scope of the current article would be necessary. Finally, we also looked at the CD163 surface marker, where we noted, similarly to IL-10 case, significance in both supplementation, donor variation as well as their interaction (***Figure 5D***). Whilst D1 seemed not to respond to any supplementation, D2 showed downregulation of CD163 for both LMW and HMW THA, and D3 small downregulation for HMW THA (***Figure 5D***). These responses were even more pronounced in biochemically induced controls (***Figure S7J-L***) for donors 2 and 3.

In the overall picture, we observed that D2 had largest capacity to respond to the supplementation of both LMW and HMW THA and that the generated MΦs appear to point towards a heterogenous state spanning some characteristics of M1 and some of M2a subtypes. We highlight that the gene expression experiments show difficulty of assessing the polarization states of PBMCs-derived MΦs supplemented with THA. Rather larger discrepancy is observed to the data obtained for the THP-1-derived MΦs, pointing out that differences in cellular origin, differentiation status as well as genetic/epigenetic factors being key in this case (with statistical significance noted across 3 tested genes for influence of donors). To further investigate the influence of THA supplementation, in the last step we evaluated cytokine expression by performing ELISAs on selected cytokines and donors.

#### 3.3.4. Cytokine expression confirmation

Secretion of TNF-α was below the limit of detection (LoD) of 15.6 pg mL^-1^ for all M0 and M2 states across all groups (control, LMW and HMW THA), and was only evident in M1 groups of all donors (***Figure 6***), confirming expected most pro-inflammatory profile for the biochemically induced groups already discussed. For D1 and D3, there were no significant differences between control, LMW or HMW conditions, whereas for D2, secretion of TNF-α was higher when supplemented with LMW and HMW THA, (p<0.01, ***Figure 6A***). This indicated a larger susceptibility of D2 to it and reaffirmed susceptibility observed in gene expression data for this donor (***Figure 5***).

**Figure 6.**
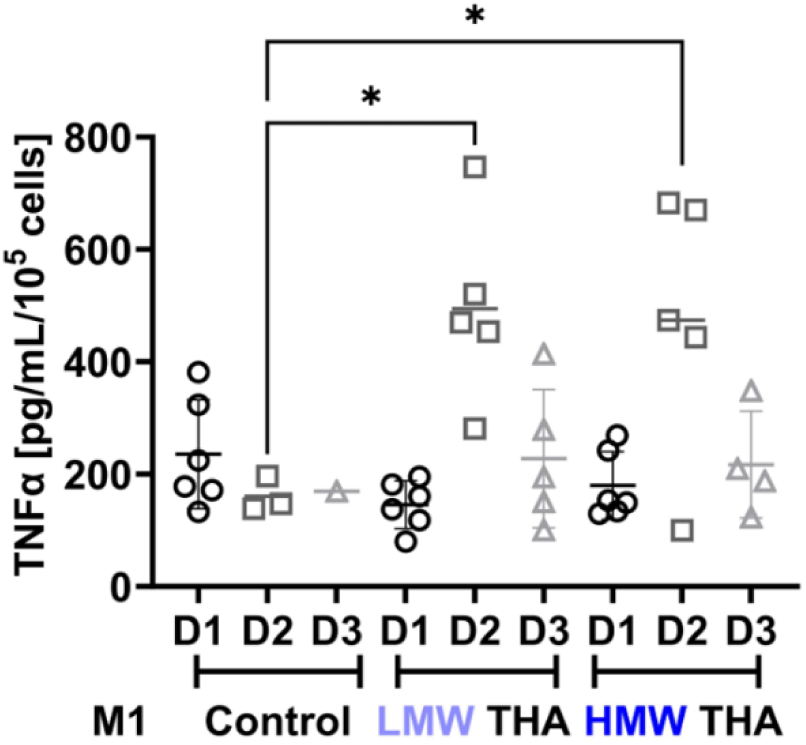
TNF-α secretion measured by ELISA for cells without any material conditioning (control) or encapsulated with 2 mg mL^-1^ LMW or HMW THA, represented for each donor (D1, D2 and D3). Data is represented only for the M1 state, pre-stimulated by the addition of 10 ng mL^-1^ TNF-α and IFN-γ. The secretion for the other two cases (M0 and M2) in any other group was below the limit of detection, represented by the dotted line. Analysis by one-way ANOVA with Šídák’s multiple comparisons test. A statistically significant results were considered for p<0.05 (* - <0.05, ** - <0.01, *** - <0.005 and **** - <0.001).

We also evaluated secretion of IL-6, IL-10, IL-1β, and CD163 for D1, and IL-10 and CD163 for D2 and D3. For none of the donors we detected any IL-10 (with the LoD of 31.3 pg mL^-1^), corroborating with the gene expression data showing no significant upregulation for any of the groups and polarization states. Combining above data for CD206 markers obtained for PBMC-derived MΦs (***Figure 4I***), this indicates that speculatively these MΦs do not tend to polarize towards any other subtype than M2a. IL-1β for D1 also has not been detected for any of the groups (with LoD = 3.91 pg mL^-1^). In D1, IL-6 was largely undetected (LoD=9.38 pg mL^-1^) for control and HMW groups (***Figure S8A***), and only low amounts were detected in LMW THA group (in all states, M0, M1 and M2). For the CD163, we have detected none for D2 (LoD=156 pg mL^-1^), and variable amounts for D1 (***Figure S8B***) and D3 (***Figure S8C***). The only statistically significant difference was observed between M1 control group and M0 supplemented with LMW THA in D1 (***Figure S8B***). On the other hand, no significant differences were observed between any polarisation states or groups in D3 (***Figure S8B***). This indicates that CD163, although often describing well the polarisation states, is strongly donor dependent for PBMCs-derived MΦs.

## 4. Conclusions

In summary, we investigated polarization tendencies of THP-1 and PBMCs-derived MΦs upon supplementation with soluble chemically tyramine-modified hyaluronan (THA) at 2 mg mL^-1^, a key biopolymer signal and building block for many biomaterials.

Flow cytometry and multiplex ELISA showed that LMW THA induces a pro-inflammatory M1-like phenotype in THP-1-derived MΦs, while HMW THA led to a mixed M1/M2 profile, suggesting a heterogeneous or transitional state. PBMCs-derived MΦs in all groups expressed CD68, confirming successful differentiation from monocytes. We also noted highest number of CD206^+^ cells for group supplemented with HMW THA, and reduction of M2 population in a group supplemented with LMW THA. Combining gene expression and ELISA data, in PBMCs-derived MΦs, both LMW and HMW THA induced transitional M1/M2 states with notable donor variability.

Differences in gene expression and proteins markers between PBMCs and THP-1-derived MΦs may be due to their cellular origin, genetic mutations and genetic/epigenetic factors, warranting further analysis for better control. Additionally, the limited number of cells obtained from PBMC donors reduced the number of tests that could be performed. The use of single THA concentration is a limitation, as concentration may modulate responses. Beyond known HA receptors interactions (like CD44 or RHAMM), other receptors and compensatory pathways may play a role, suggesting the need for future detailed mechanobiological studies. In this study we hence provide a baseline immune profile (e.g., cytokine levels, surface markers) of PBMC-derived MΦs from three different donors prior to the potential future treatment with THA-based biomaterials. Overall, LMW and HMW THA supplementation showed immunomodulatory profiles similar to non-modified HA observed in the literature, indicating that tyramine modification preserves these properties. This work provides valuable insights for applying translational THA-based biomaterials in tissue engineering.

## Supporting information

Electronic Supplementary Information

## Author Contributions

**J.K.W.**: Conceptualization, data curation, formal analysis, funding acquisition, investigation, methodology, project administration, resources, supervision, validation, visualization, writing -original draft, writing – review & editing; **E.I.B.**: Data curation, investigation, methodology, writing – review & editing; **A.J.V.**: Data curation, investigation, methodology, writing – review & editing; **M.M.**: Data curation, investigation, methodology; **M.A.**: investigation, methodology; **P.S.T.**: Data curation, methodology; **J.S.** Data curation, investigation, methodology, formal analysis, writing – review & editing; **J.T.** Conceptualization, resources, funding acquisition, supervision, writing – review & editing; **D. E.**: Conceptualization, funding acquisition, supervision, writing – review & editing; **M.D.**: Conceptualization, funding acquisition, methodology, project administration, resources, supervision, writing – review & editing.

## Acknowledgements

The authors thank Nicolas Devantay for his technical support, particularly with the THA synthesis and Dr. Ursula Menzel for her assistance with PCR plating. The graphical abstract was created with BioRender.com.

## Supplementary data

Supplementary data available: ^1^H NMR spectra of synthesized THA; bright field microscopy images; NFκB-GFP fluorescence signals measured by flow cytometry; additional imageJ analysis and immunofluorescence images, detailed RT-qPCR gene analysis and ELISA results. Supplementary data to this article can be found online at XYZ.

## Notes

All research data supporting this publication are directly available within this publication and associated supporting information.

