## Supplementary material for "Effect of molecular weight of tyramine-modified hyaluronan on polarization state of THP-1 and peripheral blood mononuclear cells-derived macrophages": Electronic Supplementary Information

[c] Independent Scholar

[d] Laboratory for Immuno Bioengineering Research and Applications, Division of Engineering, New York University Abu Dhabi, Abu Dhabi, 129188, United Arab Emirates

[e] Mines Saint-Étienne, Univ Jean Monnet, INSERM, U1059 Sainbiose, Saint-Étienne, France

[δ] Current address: Henry M. Rowan College of Engineering, Department of Chemical Engineering and Biomedical Engineering, Rowan University, Glassboro, NJ, USA

*Corresponding author J.K.W.

**NMR analysis**

For NMR analysis, a 3 w/v% of THA was dissolved in 1 mL of 0.04 w/w% hyaluronidase in $D_2O$ at 37°C and left overnight for full dissolution, before measurement on $^1H$ NMR, 300 MHz, Bruker Avance III, taken at room temperature.

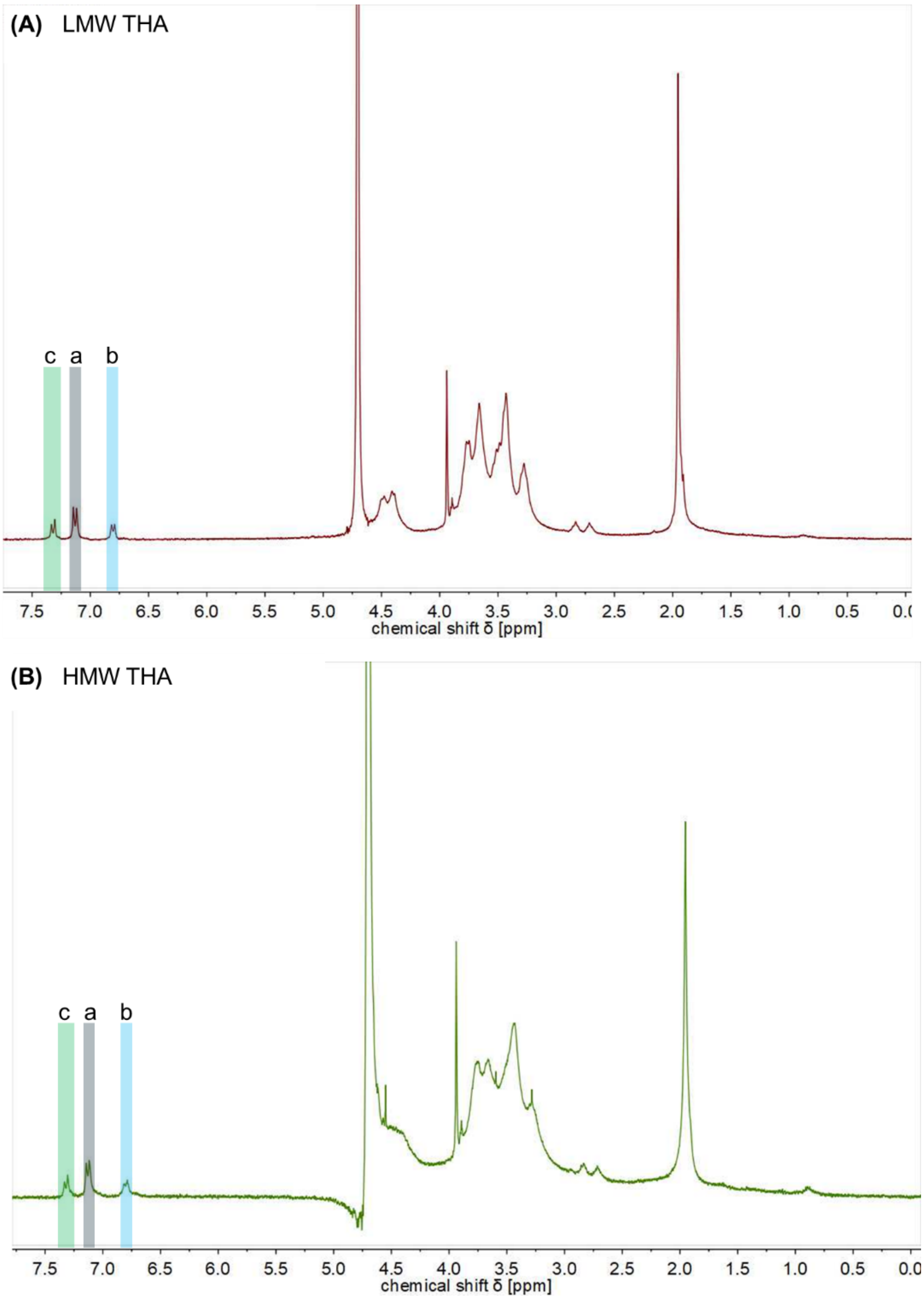


**Figure S1.** Representative $^1H$ NMR spectra of **(A)** LMW THA and **(B)** HMW THA. Highlighted regions are characteristic for the tyramine functionalization of HA and conform to the previously published spectra [1]. The presence of tyramine in the THA was verified by the resonances at 7.17–7.19 ppm (a, **grey** shading), 6.85-6.86 ppm (b, **blue** shading) and 7.36-7.38 (c, **green** shading).

**Table S1.** RNA content calculated for each donor and type of sample (averaged from 2 independent wells/replicates.

| **Donor** | **D1** | **D2** | **D3** |
|---|---|---|---|
| **Cell seeding density per well** | 1.82 x $10^5$ cells | 3.0 x $10^5$ cells | 3.0 x $10^5$ cells |
| **Sample** | **RNA content [ng], average ± SD (from 3 independent wells/replicates)** | | |
| **M0** | 561 ± 285 | 737 ± 368 | 719 ± 481 |
| **M1** | 1177 ± 505 | 824 ± 276 | 1240 ± 401 |
| **M2** | 782 ± 417 | 420 ± 67 | 706 ± 249 |
| **M0_LMW** | 1236 ± 294 | 475 ± 100 | 950 ± 174 |
| **M1_LMW** | 774 ± 695 | 436 ± 36 | 1200 ± 71 |
| **M2_LMW** | 452 ± 124 | 905 ± 241 | 1077 ± 134 |
| **M0_HMW** | 589 ± 127 | 867 ± 338 | 786 ± 236 |
| **M1_HMW** | 734 ± 327 | 555 ± 71 | 908 ± 286 |
| **M2_HMW** | 661 ± 353 | 495 ± 66 | 820 ± 437 |

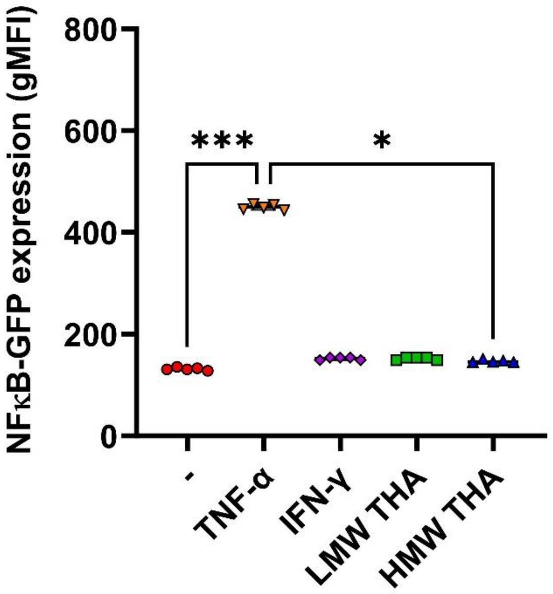


**Figure S2. (A)** NFκB-GFP fluorescence signals measured by flow cytometry. Geometric mean fluorescent signal intensities (gMFI) plotted from reported THP-1 cells directly encapsulated with 2 mg $mL^{-1}$ THA (10 times diluted from originally prepared THA hydrogels at 20 mg $mL^{-1}$ with 0.1 U $mL^{-1}$ HRP 0.15 mM $H_2O_2$) for 24 hours. 10 ng $mL^{-1}$ TNF-α was used as positive control, pure RPMI medium as negative (-) and 10 ng $mL^{-1}$ IFN-γ, used to confirm specificity of control because it does not trigger NFkB signalling pathway. Analysis by Kruskal-Wallis multiple comparisons test. A statistically significant results were considered for p<0.05 (* - <0.05, ** - <0.01, *** - <0.005 and **** - <0.001).

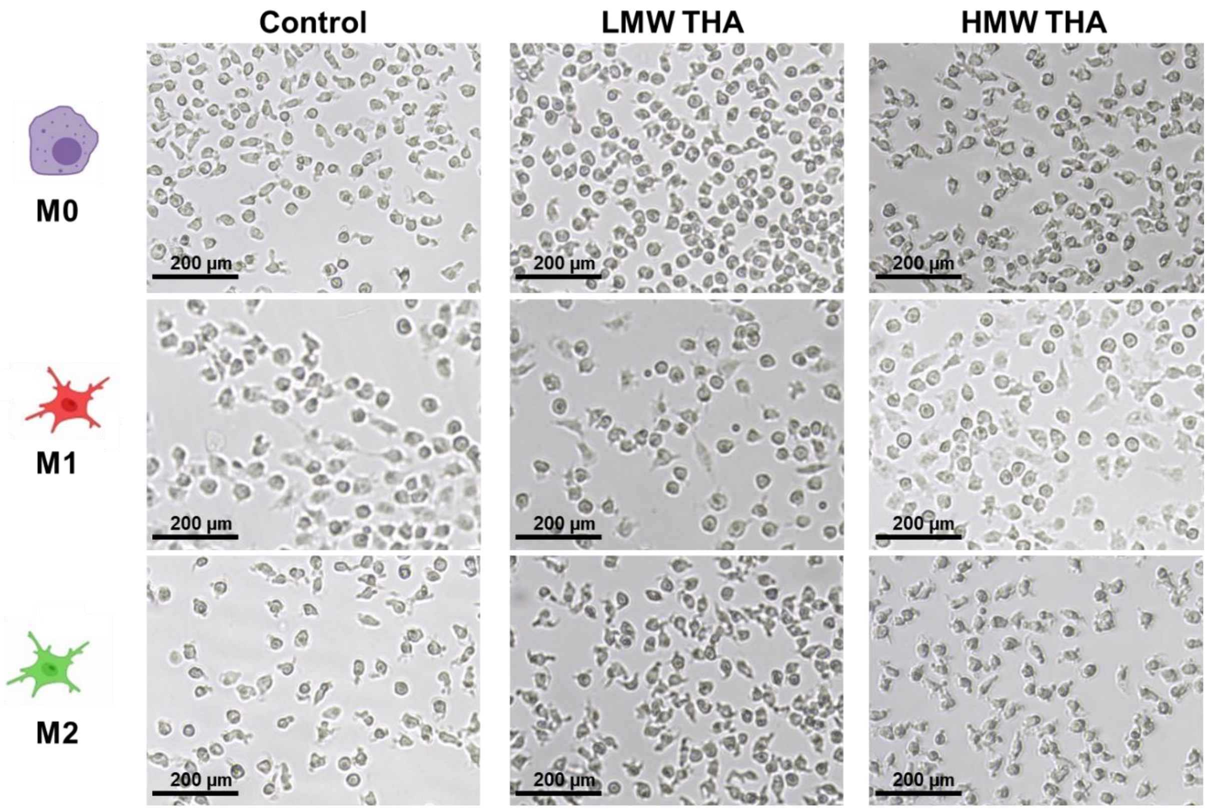


**Figure S3.** Bright field microscopical images of D1 PBMCs-derived macrophages at day 6 of the polarization protocol, just before the last 24 hours medium replenishment step.

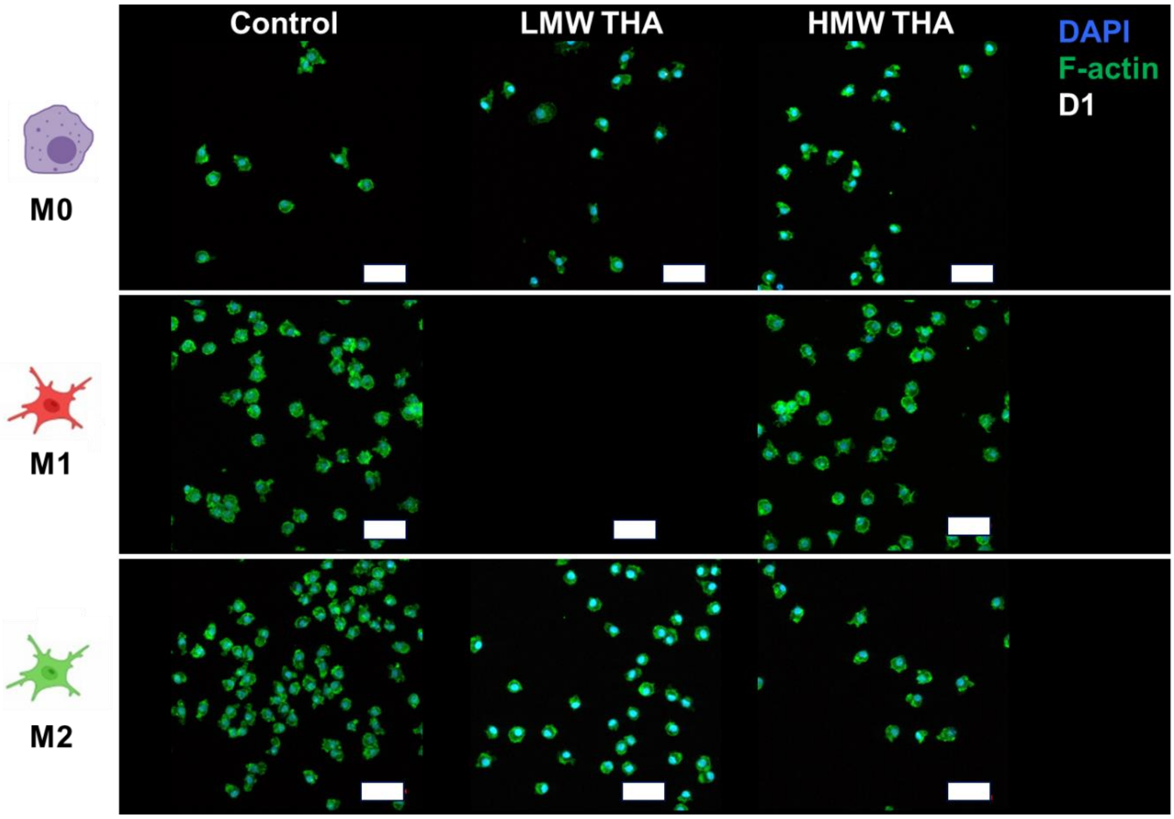


**Figure S4.** Lower magnification images of immunofluorescence staining of nucleus by DAPI (**blue**) and F-actin by Phalloidin (**green**) for all tested groups in D1 peripheral blood mononuclear cells (PBMCs)-derived macrophages. Each raw represents different polarization state (M0, M1 and M2) and columns represent a control and samples supplemented with 2mg $mL^{-1}$ LMW or HMW THA. Scale bar = 50 µm.

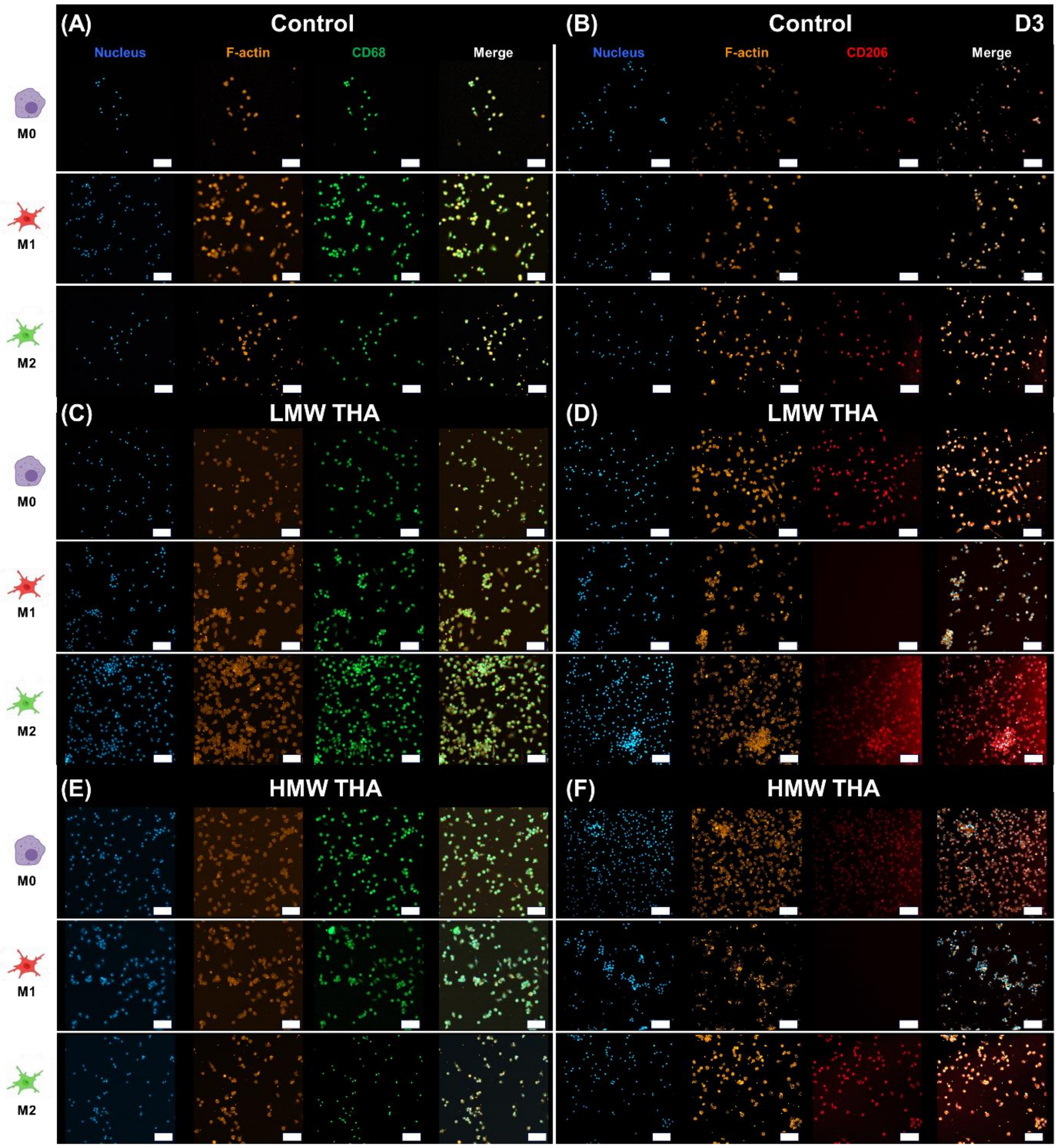


**Figure S5. Immunofluorescent characterisation of D3 peripheral blood mononuclear cells (PBMCs)-derived macrophages.** Immunofluorescence staining of nucleus by DAPI (**blue**) and F-actin by Phalloidin (**orange**) are common in all panels, for all tested groups, in donor 3 (D3) samples. Staining of CD68 (**green**) is shown in panels **(A)**, **(C)**, and **(E)**; whereas staining of CD206 (**red**) is shown in panels **(B)**, **(D)**, and **(F)**. Each row represents different polarization state (M0, M1 and M2) as indicated on the left-hand site. **(A)** and **(B)** represent panels for a control, **(C)** and **(D)** represent samples supplemented with 2mg mL$^{-1}$ LMW THA and **(E)** and **(F)** represent samples supplemented with 2mg mL$^{-1}$ HMW THA. Scale bar for all images = 100 µm.

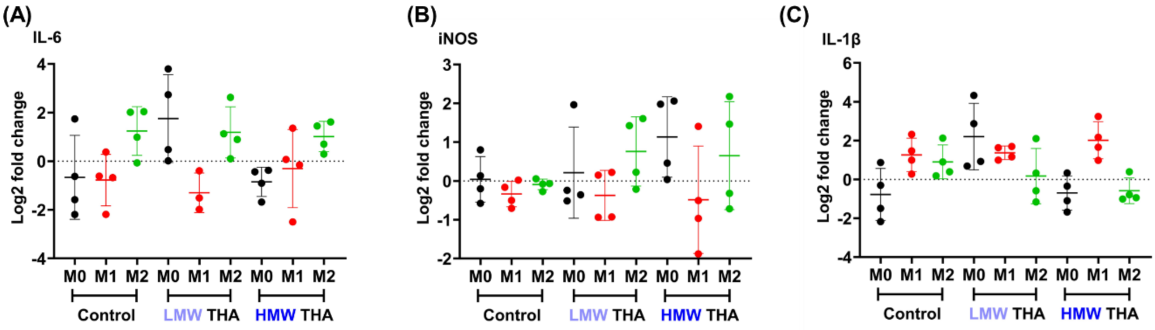


**Figure S6. (A)** IL-6, **(B)** iNOS, **(C)** IL-1β log2 fold change gene expression measured by RT-qPCR for cells without any material conditioning (control) or encapsulated with 2mg $mL^{-1}$ LMW or HMW THA, all measured for Donor 1 (D1). Data is represented for three cases: i) M0 without any additional stimulation, ii) M1 pre-stimulated by the addition of 10 ng $mL^{-1}$ TNF-α and IFN-γ and iii) M2 pre-stimulated by the addition of 10 ng $mL^{-1}$ IL-4. For easier identification, each polarization state has been marked by distinct colour on all graphs in the following way: ● - M0 ● - M1 ● - M2. Gene expression is represented as fold change compared to RPLP0 (a housekeeping gene) of the M0 control group. Data represent mean ± SD of 4 independent experiments, for D1. Analysis by one-way ANOVA with Kruskal-Wallis multiple comparisons test was done, however no statistical significance was obtained for any of the presented data.

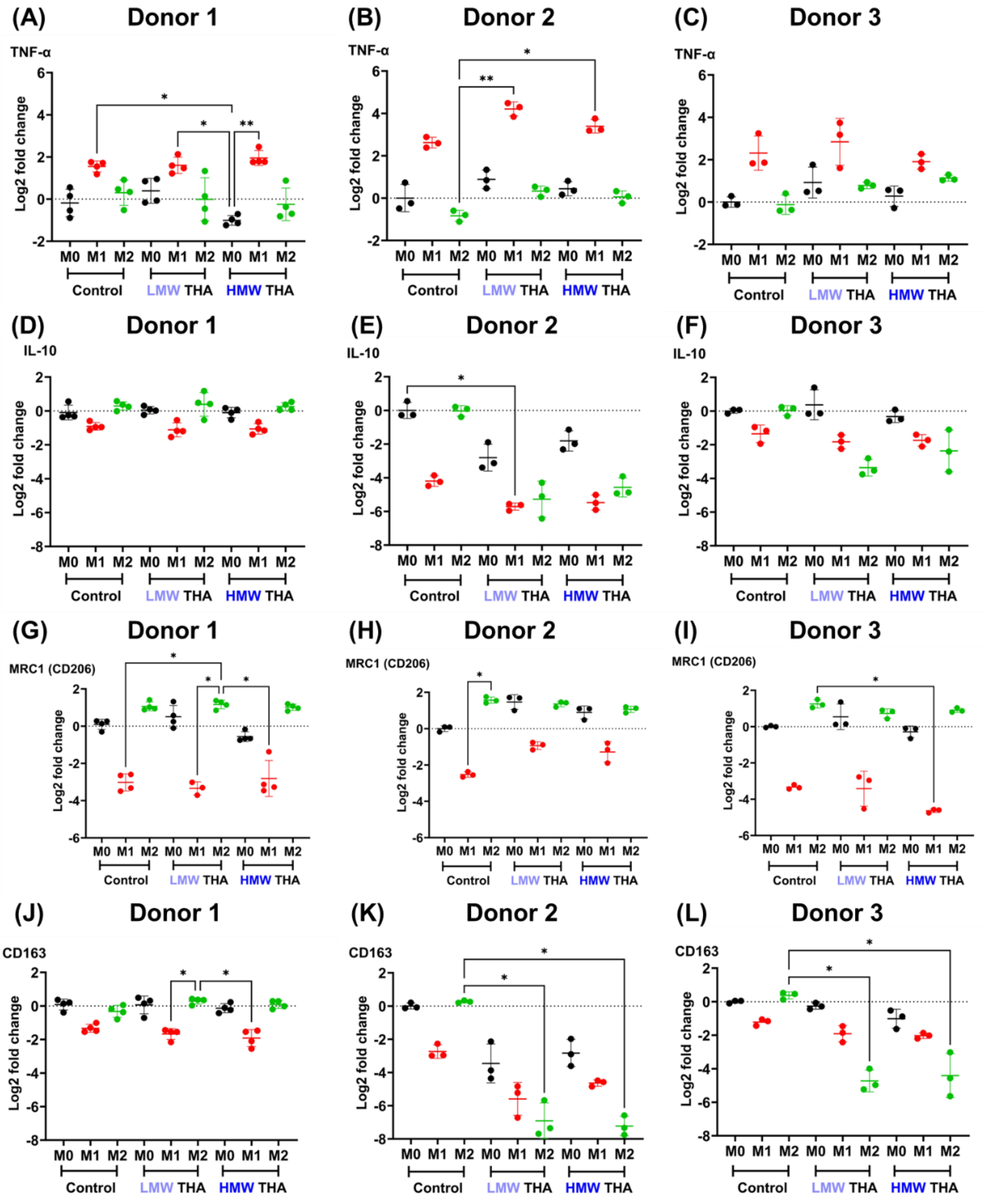


**Figure S7.** Log2 fold change gene expression of **(A-C)** TNF-α, **(D-F)** IL-10, **(G-I)** MRC1 (CD206), **(J-L)** CD163, measured by RT-qPCR for cells without any material conditioning (control) or encapsulated with 2mg $mL^{-1}$ LMW or HMW THA. Data is represented for three cases: i) M0 without any additional stimulation, ii) M1 pre-stimulated by the addition of 10 ng $mL^{-1}$ TNF-α and IFN-γ and iii) M2 pre-stimulated by the addition of 10 ng $mL^{-1}$ IL-4. For each gene, each panel presents data from individual donor, with **(A)**, **(D)**, **(G)**, **(J)** for D1, **(B)**, **(E)**, **(H)**, **(K)** for D2 and **(C)**, **(F)**, **(I)**, **(L)** for D3. For easier identification, each polarization state has been marked by distinct colour on all graphs in the following way: ● - M0 ● - M1 ● - M2. Gene expression is represented as fold change compared to RPLP0 (a housekeeping gene) of the M0 control group. Data represent n=4 for D1 and n=3 for D2 and D3 of individual experiments. Analysis by one-way ANOVA with Kruskal-Wallis multiple comparisons test was done. A statistically significant results were considered for $p<0.05$ (* - <0.05, ** - <0.01, *** - <0.005 and **** - <0.001).

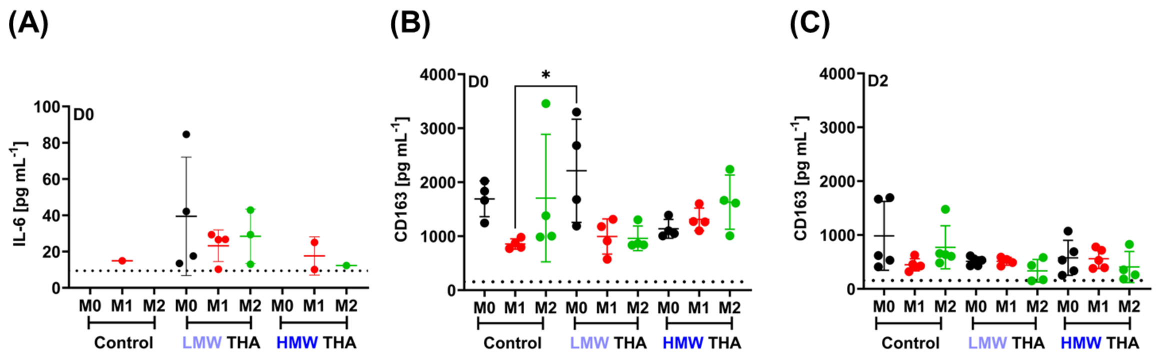


**Figure S8. (A)** IL-6 for D1, **(B)** CD163 for D1 and **(C)** CD163 for D3, secretion measured by ELISA for cells without any material conditioning (control) or encapsulated with 2mg $mL^{-1}$ LMW or HMW THA. Data is represented for three cases: i) M0 without any additional stimulation, ii) M1 pre-stimulated by the addition of 10 ng $mL^{-1}$ TNF-α and IFN-γ and iii) M2 pre-stimulated by the addition of 10 ng $mL^{-1}$ IL-4. For easier identification, each polarization state has been marked by distinct colour on all graphs in the following way: ● - M0 ● - M1 ● - M2. In each case, dotted line represents the limit of detection. Analysis by one-way ANOVA with Šídák's multiple comparisons test. A statistically significant results were considered for $p<0.05$ (* - <0.05, ** - <0.01, *** - <0.005 and **** - <0.001).
